# Individual traits shape hemispheric vlPFC sensitization to Cyberball exclusion: Evidence from single-trial fNIRS analyses

**DOI:** 10.64898/2026.06.05.730448

**Authors:** Cailee M. Nelson, Xiaoxue Fu, Laura M. Morett, Caitlin M. Hudac

## Abstract

Ostracism (i.e., social exclusion) threatens social security and individuals who are hypersensitive to it may face long-term negative mental health consequences. Clarifying how self-reported distress and neural responses to ostracism unfold moment-by-moment and across varying levels of individual traits may help better understand what contributes to hypersensitivity. To this objective, the current study employed a functional near-infrared spectroscopy (fNIRS) adapted Cyberball task and single-trial analytic techniques in a sample of 53 college students (aged 18-22 years). Results confirmed that Cyberball induced ostracism increased self-reported distress and neural activity in the ventrolateral prefrontal cortex (vlPFC), a region associated with emotion regulation. Self-reported distress varied with differences in social anxiety and need for belonging. Moreover, vlPFC activity sensitized across exclusion-specific trials and this neural sensitization was differentially modulated by social anxiety, need for belonging, and personality growth mindset. The current study highlights the value of single-trial approaches for capturing the nuanced temporal dynamics of ostracism responses. Additionally, it underscores the importance of examining how individual differences shape immediate responses to ostracism to better understand their association with downstream, longer-term consequences.

## 1. Introduction

An innate human need for belonging drives the formation and maintenance of social connections that are necessary for physical and psychological well-being (Baumeister & Leary, 1995). Ostracism, or the exclusion and ignoring of an individual, threatens an individual’s social security by thwarting one’s sense of belonging. Feeling distress after experiencing ostracism is evolutionarily advantageous as it alerts individuals to real social threats (Hales et al., 2016); yet, hypersensitivity to ostracism increases the likelihood of an individual developing mental disorders like depression and anxiety that may perpetuate ostracism (Albath et al., 2025; Büttner & Greifeneder, 2024; Reinhard et al., 2020; Rudert et al., 2021).

Understanding how psychological and neurological responses to ostracism unfold over time is key to explaining how hypersensitivity develops. A leading theory (Williams, 2009) outlines three stages that occur in response to ostracism (i.e., social need-threat): 1) an immediate *reflexive* stage where increases in psychological and biological indicators of social pain are universal, 2) a coping *reflective* stage where individuals attempt to understand and assign meaning to the ostracism, and, if ostracism persists, 3) a *resignation* stage marked by long-term negative mental health outcomes. Although mapping these temporal shifts is crucial for capturing ostracism’s full impact, prolonged ostracism is notoriously hard to model and measure. Short-term responses are easier to document, yet emerging models suggest the temporal dynamics are far more complex than once assumed and deserve further investigation (Riva et al., 2025).

Cyberball, a virtual ball-tossing game that manipulates social inclusion and ostracism (i.e., social exclusion; Williams et al., 2000), is commonly employed to better understand psychological and neural measures of distress that occur in the short-term (for meta-analyses see Cacioppo et al., 2013 and Hartgerink et al., 2015). Immediately following ostracism manipulated within the game, participants of all ages report increased threats to four fundamental social needs: belonging, sense of control, self-esteem, and beliefs of a meaningful existence (Hartgerink et al., 2015). In addition to these social threats, ostracism heightens neural reactivity in brain regions associated with conflict monitoring and emotion regulation (e.g., anterior cingulate cortex [ACC], ventrolateral prefrontal cortex [vlPFC], dorsolateral prefrontal cortex [dlPFC]; Eisenberger et al., 2003; Lieberman, 2007; Masten et al., 2011; Onoda et al., 2009).

The vlPFC is crucial for emotion regulation through response selection and inhibition (Ochsner et al., 2012), making it a critical region for studying neural sensitivity to ostracism. However, the specific functional roles of left versus right vlPFC during emotion regulation remain highly debated (Cheng et al., 2022). Ostracism research consistently reports activation in both hemispheres during social exclusion (Bolling et al., 2011; Eisenberger et al., 2003; Masten et al., 2009; Onoda et al., 2009, 2010), and transcranial magnetic stimulation (TMS) shows that stimulating either hemisphere impairs emotion regulation—though likely through different mechanisms (Cheng et al., 2022; Li et al., 2022). For example, disrupting left vlPFC reduces verbal fluency during oral reporting of emotion regulation strategies, implicating its role in language-based emotion regulation strategy generation (Cheng et al., 2022). Conversely, disrupting the right vlPFC hinders reappraisal of negative emotions, suggesting an association with inhibitory control during emotion regulation (Cheng et al., 2022). This pattern is further clarified by evidence demonstrating greater left vlPFC activity for social exclusion paired with emotionally supportive messages (Onoda et al., 2009) and negative associations between right vlPFC activity, self-reported distress, and ACC activity (Eisenberger et al., 2003; Masten et al., 2009).

Early evidence suggested immediate responses to ostracism are largely unaffected by individual differences like participant gender (Williams & Sommer, 1997), cultural dimensions (e.g., individualism/collectivism; Smith & Williams, 2004), and social anxiety (Zadro et al., 2006). Yet growing evidence challenges this view, suggesting that traits such as features of personality disorder and fears of social threat modulate the intensity of self-reported distress (Boyes & French, 2009; Riva et al., 2014; Wirth et al., 2010). Neuroimaging research provides additional support for these behavioral findings. For example, individuals with low self-esteem demonstrate more connectivity between right vlPFC and regions of the ACC compared to those with high self-esteem, indicating reduced ability to regulate social pain generated by the ACC (Onoda et al., 2010). Individuals with lower rejection sensitivity demonstrate greater left prefrontal cortex activity when viewing images of rejection compared to individuals with greater rejection sensitivity (Kross et al., 2007). Less tolerance for uncertainty (i.e., excessive worrying of or avoiding uncertain situations) is associated with increased levels of trait aggression via reduced vlPFC activity during social exclusion (Gorka et al., 2018). Finally, although social anxiety does not influence DLPFC activity during social exclusion alone, it is associated with heightened DLPFC activity when receiving emotional support during social exclusion (Nishiyama et al., 2015). Together, these findings underscore that short-term hypersensitivity to ostracism can be influenced by individual differences. Still, more precise characterization of how responses to ostracism unfold in real time (i.e., across the course of the experiment) is needed to clarify how different levels of sensitization relate to long-term mental health outcomes.

### 1.1. Current objectives

In addition to fMRI research, more recent functional near-infrared spectroscopy (fNIRS) studies have used Cyberball to examine neural correlates of ostracism across regions in prefrontal cortex associated with emotion regulation (Dou et al., 2020; Jiao et al., 2024; Lian et al., 2025; Song et al., 2024). Given that this body of work is much smaller, the first aim of the current study was to examine classic differences in self-reported social threats and neural sensitivity during the ostracism block compared to the fair play (i.e., solely inclusionary) block in an adapted fNIRS Cyberball task. In line with the previous literature, we hypothesized that ostracism in comparison to fair play would elicit increases in self-reported social threats and vlPFC activity (Eisenberger et al., 2003; Hartgerink et al., 2015; Lieberman, 2007; Masten et al., 2011; Onoda et al., 2009). Due to conflicting findings of left versus right vlPFC activity (Bolling et al., 2011; Eisenberger et al., 2003; Masten et al., 2009, 2011; Onoda et al., 2009, 2010) we opted to include hemisphere to evaluate and report data-driven findings. Finally, as vlPFC is associated with emotion regulation, we hypothesized that increased self-reported social threats would be related to decreased vlPFC activity (Eisenberger et al., 2003; Masten et al., 2009).

Second, modern neuroscience approaches use single-trial analyses to better understand real-world interactions (Stokes & Spaak, 2016). Despite a well-known theoretical model that emphasizes temporal dynamics of ostracism responses (Williams, 2009) and the increasingly apparent social exclusion that occurs during the Cyberball ostracism block, we still lack detailed understanding of the moment-to-moment trial-level changes that occur within this paradigm. Cyberball studies often manipulate fair play blocks by having other “players” throw the ball to each other and the participant an equal amount of time across the block (i.e., solely inclusionary), in contrast to the ostracism block that begins with a short inclusion phase and ends with a longer exclusion phase (i.e., partial ostracism, ambiguous ostracism; Boyes & French, 2009; Eisenberger et al., 2003; Masten et al., 2009, 2011; Onoda et al., 2009; Riva et al., 2014; van Noordt et al., 2015; Williams et al., 2000). However, it remains unclear if neural activity changes within the time spent being excluded in the ostracism block. For example, it is reasonable to assume that participants may not initially realize that they are being excluded until later in the task and, thus, could show neural sensitization over time that may also be a better predictor of downstream mental health consequences. As such, the second aim of this study was to use a single-trial analysis to evaluate changes in vlPFC activity over time spent in each phase of the Cyberball task. We hypothesized that vlPFC activity would increase over trials within the exclusion phase of the ostracism block as participants become more aware that they are being excluded.

Lastly, as argued by Riva and colleagues (2025), there is growing evidence that individual differences are increasingly more apparent, particularly related to short-term psychological and neurological processing of ostracism. As such, the current study set out to investigate how individual differences modulated self-reported social threats, neural sensitivity between blocks, and sensitization during the exclusion phase of the ostracism block. Here, we opted to explore individual differences in social anxiety, need for belonging, and personality growth mindset because of their known associations with emotion regulation and sensitivity to ostracism. For example, individuals with social anxiety use different emotion regulation strategies (Dryman & Heimberg, 2018) and demonstrate significantly greater vlPFC activity during reappraisal of negative emotions (Campbell-Sills et al., 2011). Additionally, need for belonging has been associated with negative social emotions, neuroticism, and rejection sensitivity (Leary et al., 2013), suggesting that individuals with higher need for belonging may be more sensitive to instances of social exclusion. In contrast, individuals with a personality growth mindset (i.e., the belief that personality features are malleable; Yeager & Dweck, 2012) when faced with exclusion or bullying are more likely to educate their peers about why these behaviors are harmful rather than respond with retaliation (Yeager et al., 2011; Yeager & Miu, 2011). Taken together, we expected that more social anxiety, need for belonging, and fixed mindsets of personality would demonstrate heightened vlPFC activity during the ostracism block and increased sensitization in vlPFC activity during the exclusion phase compared to those with less social anxiety, need for belonging, and growth mindsets of personality. Like the first aim, we include vlPFC hemisphere to more fully interrogate data-driven effects, given mixed findings of distinct hemispheric roles.

## 2. Method

### 2.1. Participants

Sixty-one college students (aged 18 to 22 years) participated in the current study. Participants were recruited from research pools and local autism databases from the University of Alabama. All procedures for this study were performed in compliance with institutional guidelines approved by the university’s internal review board. Written informed consent was obtained from all participants prior to participation. Due to the deceptive nature of Cyberball, all participants were debriefed following experimental sessions, informed of the true nature of the study, and asked if they were aware they were being deceived. About 31% of the sample (19 of 61) indicated that at some point they were aware that they were being deceived. Due to technical issues during fNIRS data acquisition and extremely noisy data, eight participants were not included in the final analyses. Demographic information for the final sample (*n* = 53) can be found in Table 1.

**Table 1.** Demographic information for study sample.

|  | *n (%)* |  |  |
| --- | --- | --- | --- |
| **Gender** |  |  |  |
| Female | 34 (64.81) |  |  |
| Male | 19 (35.18) |  |  |
| **Ethnicity** |  |  |  |
| Hispanic/Latine | 2 (3.70) |  |  |
| Not Hispanic/Latine | 51 (96.30) |  |  |
| **Race** |  |  |  |
| American Indian/Alaska Native | 2 (3.70) |  |  |
| Black | 3 (7.41) |  |  |
| White | 48 (88.89) |  |  |
| **Neurodiverse Conditions** |  |  |  |
| Autism | 2 (3.70) |  |  |
| Sensory Impairments (e.g., vision<br>or hearing loss) | 1 (1.89)<br>1 (1.89) |  |  |
| Mobility Impairments | 6 (11.32) |  |  |
| Learning Disability (e.g., ADHD,<br>dyslexia) | 4 (7.55)<br>1 (1.89) |  |  |
| Mental Health Disorder |  |  |  |
| Other Condition-Not Specified |  |  |  |
|  | *M (SD)* | *Possible Range* | *Sample Range* |
| **Age** | 19.70 (0.97) | 18-25 | 18-22 |
| **Liebowitz Social Anxiety Scale (LSAS)** | 45.85 (24.54) | 0-144 | 4-117 |
| **Need to Belong Scale (NTBS)** | 35.45 (6.49) | 10-50 | 19-49 |
| **Implicit Theories of Personality (ITP)** | 10.51 (3.95) | 5-30 | 5-21 |

**Note.** For age and individual difference measure scores we report the sample's mean (*M*) and standard deviation (*SD*), possible range, and the sample's actual range.

### 2.2. Measures to account for individual differences

#### 2.2.1. Need-Threat Survey

The Need-Threat Survey (NTS; Williams, 2009) assessed participants’ awareness of exclusion and self-reported social threats after each Cyberball block. This survey consisted of 20 items targeting perceived social threats to four fundamental needs (i.e., belonging, sense of control, beliefs of a meaningful existence, self-esteem), eight mood items (e.g., “My mood was good”, “My mood was unfriendly”), and three items addressing the manipulation of exclusion (i.e., the percentage of tosses they received, how excluded they felt). All items except perceived toss percentage were rated on a five-point Likert scale (1 = not at all, 5 = extremely often). Likert-style items were summed across items in seven separate subdomains: feelings of exclusion, positive affect, negative affect, and social threats to belonging, control, meaningful existence, and self-esteem. Higher scores indicate greater feelings of exclusion, affect, and social threats.

#### 2.2.2. Implicit Theories of Personality Scale

The implicit theories of personality scale (ITP; (Yeager et al., 2011) measures implicit theories (i.e., mindset) about how people’s—specifically, bullies’, victims’, winners’ and losers’—personality traits can change. Individuals rated five items on a six-point scale where one indicates “strongly disagree” and six indicates “strongly agree.” Items were summed for a total score where lower scores indicate a growth mindset of personality.

#### 2.2.3. Liebowitz Social Anxiety Scale

The Liebowitz Social Anxiety Scale (LSAS; Liebowitz, 1987) was used to measure social anxiety features in the sample. This scale consists of 24 items that aim to measure how social anxiety plays a role in a person’s life across various situations. Participants were asked to rate each item based on two questions: how anxious they feel about the situation and how often they avoid the situation. Both sets of questions were rated on a zero to three scale where zero represents no anxiety or no avoidance and three represents severe anxiety and severe avoidance. Participants’ responses were summed to create a total social anxiety scale where scores ranging from 0 to 29 represent no social anxiety, 30-49 mild social anxiety, 50 to 64 moderate social anxiety, 65 to 79 marked social anxiety, 80 to 94 severe social anxiety, and 95 or above very severe social anxiety.

#### 2.2.4. Need to Belong Scale

The Need to Belong Scale (NTBS) is a well-known scale that measures desire for acceptance and belonging (Leary et al., 2013). Participants rated 10 items on a five-point Likert scale where one indicates “strongly disagree” and five indicates “strongly agree.” Items one, three, and seven were reverse-scored and then all items were summed to create a final score. Higher scores indicate a stronger need for belonging.

### 2.3. Cyberball task

To manipulate ostracism, we used the traditional triadic ball-throwing Cyberball task (Williams et al., 2000) with modified timing. In this task, participants completed two blocks that proceeded in the same exact order for every participant (see Figure 1A). At the start of each block, participants were shown cartoon images that they were told represented two other “real” players and themselves. To help ensure participants believed they were playing with real people, the other players were given names and participants input their own name which appeared below their character. At the start of each block, a screen with the phrase “Please wait while other players join…” was presented for 10 s. Additionally, at the end of each block, participants completed survey measures (i.e., Need-Threat Survey).

**Figure 1.**
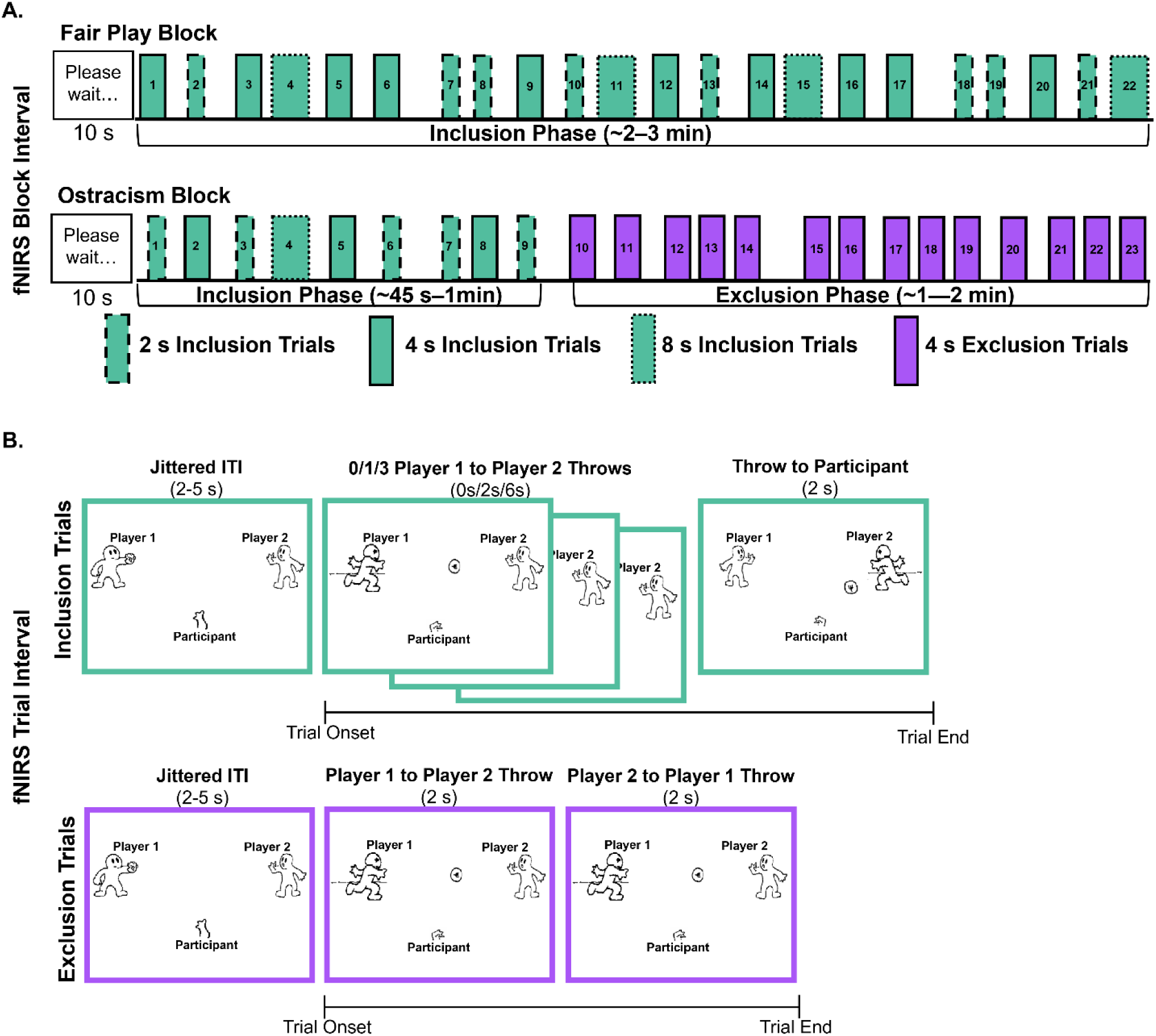
fNIRS Cyberball design. **<u>Panel A</u>** represents block- and phase-level timing. Each block started with a 10-second screen indicating that the game was looking for players to aide the ruse that participants were playing with real people. Numbers within each rectangle indicate trial order within each block. Turquoise represents inclusion trials and purple indicates exclusion trials. Because inclusion trials varied based upon number of throws between all players, trial lengths are marked by bordered lines: 2 second trials (dashed), 4 second trials (solid), and 8 second trials (dotted). **<u>Panel B</u>** illustrates timing for inclusion and exclusion trials. The top turquoise panels represent the timing of all possible inclusion trials. The bottom purple panels represent the timing for exclusion trials.

We opted for an event-related Cyberball design where participants saw a total of 45 trials (108 throws total) over two blocks: fair play and ostracism. All trials were predetermined by the research team, like other event-related Cyberball designs (see van Noordt et al., 2015). The fair play block consisted of 66 throws in which the other players threw to one another 22 times and the participant threw back to the other players 22 times (i.e., 22 inclusion trials; see Figure 1A). The ostracism block consisted of a total of 52 throws which were separated by two phases to maintain deception. At the start of the block (i.e., the inclusion phase), participants threw the ball nine times across 24 total throws (i.e., nine inclusion trials). In the second part of the block (i.e., the exclusion phase), the participants were fully excluded for 28 throws (i.e., 14 exclusion trials). To mitigate carryover effects from the ostracism block, participants always started with the fair play block (Eisenberger et al., 2003; Masten et al., 2011; van Noordt et al., 2015). Individual trial timing for trials used for analyses can be found in Figure 1B. Each trial was preceded by a jittered intertrial interval (2-5 s). Throws were always 2 s long, but total trial duration could vary in length based on the number of throws seen. For inclusion trials, participants viewed zero, one, or three throws between the other players before being thrown the ball. In this way, inclusion trials were either 2, 4, or 8 s long. Exclusion trials only consisted of two throws and were always 4 s long. fNIRS data were time-locked to the beginning of the sequence of throws in a trial (see Figure 1B).

### 2.4. fNIRS data collection

During the Cyberball task, fNIRS data was recorded using a NIRScout System (NIRx Medical Technologies, LLC, Los Angeles, CA, USA) employing near-infrared light at 760- and 850 nm wavelengths and a 7.81 Hz sampling frequency. Eight light sources and four detectors were placed over the scalp using a fNIRS cap in accordance with the standard 10-20 international system (Figure 2; Homan et al., 1987) and source-detector distance was preserved at an average of 31.9 mm. The fNIRS cap was positioned using standard procedures, centered equidistantly between the nasion-inion and preauricular landmarks. Using the fNIRS Optodes’ Location Decider toolbox (fOLD; Zimeo Morais et al., 2018), eight sources were positioned at FC5, FC6, F7, F3, F4, F8, AF7, and AF8 and the four detectors were positioned at FC3, FC4, F5, and F6 to create 12 channels corresponding to the most suitable Brodmann areas (BA) for vlPFC (e.g., 44, 45; O’Reilly, 2010). Changes in the concentrations of HbO and HbR were recorded during Cyberball using NIRStar Acquisition Software (NIRx Medical Technologies LLC, Glen Head, NY, USA).

**Figure 2.**
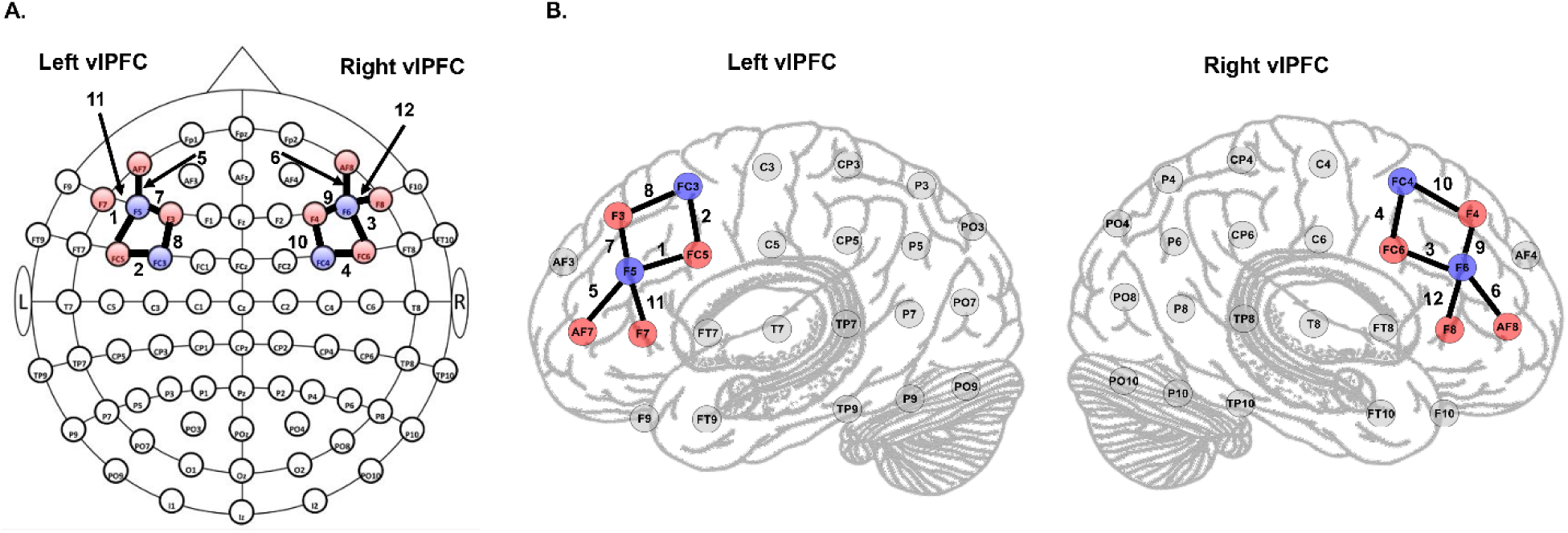
Optode locations in accordance with 10-20 international system. Across both panels, red circles indicate sources and blue circles indicate detectors. Solid black lines indicate channels. **<u>Panel A</u>** represents an overhead view of the montage with channel labels numbered to the side of channel lines. **<u>Panel B</u>** represents a lateral view of each hemisphere with channel labels numbered to the side of channel lines.

### 2.5. fNIRS data preprocessing

fNIRS data were preprocessed using Homer3 (v1.86.0; Huppert et al., 2009). Channels with poor signal-to-noise ratio in the raw signal were pruned from analyses (dRange = 3e-02—2.0e00, SNRthresh = 5, SDrange = 0.0—50.0) before the raw signal was converted to optical density. Across the entire sample, only 2.04% of channels were removed prior to analyses. Motion correction was applied using the spline interpolation with Savitzky-Golay filtering (p = 0.99, FrameSize_sec = 10; Jahani et al., 2018). We then removed high-frequency noise by applying a low-pass filter (0.50) and converted the optical density to oxygenated hemoglobin (HbO) and deoxygenated hemoglobin (HbR) concentration changes using the modified Beer-Lambert Law (ppf = 1). Pathlength correction was not applied and, thus, changes in signal were extracted as concentration changes multiplied by the mean path length (µM*cm; Yücel et al., 2021). To estimate the hemodynamic response function (HRF) from – 2 to 10 s, we opted to use a general linear model (GLM) with the least-squares method for estimating the weights of consecutive Gaussian functions (glmSolveMethod = 1, idxBasis = 1, paramdriftOrder = 3) as it allows for flexibility in estimating the HRF and is optimal for single-trial analyses (von Lühmann et al., 2020) To remove systemic physiological noise, the average of all channels was included as a regressor in the GLM (flagNuissanceRMethod = 2; Klein et al., 2022).

To better evaluate changes in neural sensitivity over time, we opted to extract baseline-corrected (-2 to 0 s) HbO values (i.e., ΔHbO) from the GLM modeled HRF time-series at the single-trial level for each channel. As such, we were able to delineate three separate phases: fair play block inclusion phase, ostracism block inclusion phase, ostracism block exclusion phase. In line with other fNIRS research (see Su et al., 2023 and Dou et al., 2020), we then normalized HbO as single trial-level *z*-scores for every individual channel using the following equation:

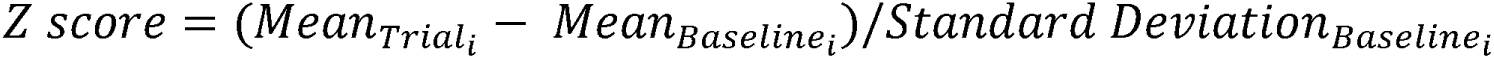

Trials with *z*-scores greater or less than ± 2 standard deviations of the average trial *z*-score were identified as outliers and removed from further analyses. Across all participants, an average of 98.75 trials (8.10%) and 99 trials (8.49%) were removed from each channel for the ostracism block and fair play block respectively.

### 2.6. Preliminary analyses and analytic plan

All analyses were conducted in R (version 4.3.3) using linear mixed-effects models that were computed using restricted maximum likelihood with Nelder-Mead optimization via the *lme4* package (Bates et al., 2015). False-discovery rate (FDR) correction (Benjamini & Hochberg, 1995) was applied for multiple comparisons for all *post-hoc* testing. As a base, all models were fit with a random intercept for each person to account for repeated measures. Additionally, when self-reported social threats (e.g., belonging, control, meaningful existence, self-esteem) and individual difference measure scores (e.g., social anxiety, need for belonging, mindset) were used as predictors in a model, values were mean centered to aid in interpretation. Models varied in their fixed effects and outcomes which are discussed in further detail below.

#### 2.6.1. Self-reported distress following Cyberball manipulation

Consistent with prior Cyberball research, all Need-Threat Survey subdomains were used to verify the ostracism manipulation. If successfully manipulated, the ostracism block should elicit greater feelings of exclusion, reduced positive affect, increased negative affect, and heightened social threats relative to the fair play block (Gorka et al., 2018; Jiao et al., 2024; Masten et al., 2011; Wirth et al., 2010). To test this, we ran separate models for each of the nine variables: estimated percent of throws received, feelings of exclusion, positive affect, negative affect, and social threats to belonging, self-esteem, meaningful existence, and control:

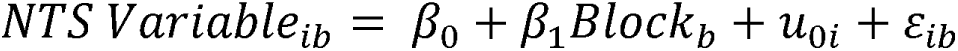

Additionally, we explored whether self-reported social threats (i.e., belonging, control, self-esteem, and meaningful existence) were influenced by individual differences. Each mean-centered individual difference variable (social anxiety, need for belonging, mindset) was entered as a main effect and as an interaction with block in separate models for each social threat outcome (i.e., three models per outcome; see Supplemental Materials 1.1 for full R syntax).

#### 2.6.2. Changes in vlPFC activity across blocks

Given that inclusion trials could vary in duration across each block based upon the number of throws between the players (1, 2, or 4 throws before re-including the participant), we conducted preliminary sensitivity analyses to determine if there were meaningful differences between inclusion trials within and across each block. We found that inclusion trials of all lengths were more negative in the ostracism block inclusion phase compared to the fair play inclusion phase (*p* < 0.0326, Supplemental Table 1) and in most cases followed a graded pattern such that longer trial types were more negative (see Supplemental Table 2). Although the 4 s trials in the fair play block inclusion phase were more negative than the other trial types within that same block, these trial types had the smallest difference between blocks across all channels. Additionally, these trial types were the most like exclusion trials (i.e., 2 throws across 4 s). Thus, we opted to target our analyses on all 4 s inclusion (*n* = 13) and exclusion trials (*n* = 14).

In line with classic Cyberball analyses, we first evaluated differences in channel-level *z*-scored ΔHbO between the fair play and ostracism blocks across the left and right vlPFC. To do this, we fit a model that included fixed and interacting effects of block (fair play, ostracism) and hemisphere (left, right). The model also included random intercepts for each participant in each hemisphere and a random slope of block to account for individual variability:

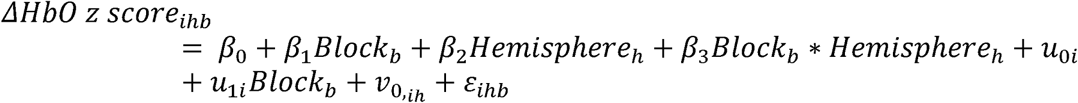

We also evaluated how self-reported social threats and individual differences modulated block effects by fitting three similar models based on the individual difference measure of interest. In this way each model included fixed and interacting effects for the mean-centered individual difference measure scores (e.g., social anxiety, need for belonging, mindset), each mean-centered social threat (e.g., belonging, control, meaningful existence, self-esteem), block, and hemisphere (see Supplemental Materials 1.2 for R syntax).

#### 2.6.3. Evaluating vlPFC sensitization to exclusion

To account for differences in inclusion trials across blocks and to investigate how channel-level *z*-scored ΔHbO became sensitized over inclusion and exclusion trials, we fit a model with phase (fair play block inclusion phase, ostracism block inclusion phase, ostracism block exclusion phase), hemisphere (left, right), and trial number (continuous) as main and interacting effects. The model also included random intercepts for each participant in each hemisphere and a random slope of phase to account for individual variability:

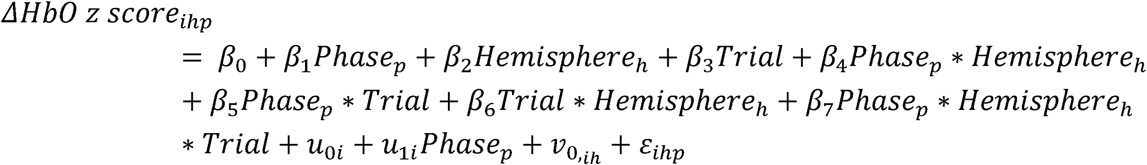

#### 2.6.4. Exploring the effect of individual differences on vlPFC sensitization to exclusion

Finally, in line with our hypotheses that individual differences would influence sensitization of vlPFC to exclusion, we explored the effects of participant characteristics on channel-level *z*-scored ΔHbO solely across trials within the ostracism block exclusion phase. To do this, we fit three models each with fixed and interacting effects for the mean-centered individual difference measure of interest (social anxiety, need for belonging, mindset), hemisphere (left, right), and trial number (continuous). Given that this was exploratory and only evaluated exclusion specific trials, we only included random intercepts for each participant in each hemisphere (see Supplemental Materials 1.3 for R syntax).

## 3. Results

### 3.1. Manipulation check

Analyses confirmed that the ostracism block successfully manipulated feelings of exclusion across all Need-Threat Survey variables, *p* < 0.0001 (Supplemental Table 3). Specifically, these analyses revealed that after the ostracism block participants estimated lower percentages of receiving the ball, felt more excluded, and reported greater social threats to belonging, control, meaningful existence and self-esteem compared to the fair play block.

### 3.2. Evaluating the influence of individual differences on self-reported distress following Cyberball

Means (*M*) and standard deviations (*SD*) for each individual difference measure are reported in Table 1. Between the individual difference measures, there were no significant Pearson correlations after FDR correction (*p* = 0.77). Each linear mixed-effects model’s results can be found in Table 2. All social threats were significantly predicted by social anxiety, such that higher levels of social anxiety were associated with higher levels of perceived threat, in general. Post-hoc analyses revealed that these estimates were only significantly different from zero for belonging, control, and self-esteem threats (*p* < 0.014; Figure 3A–3C). Need for belonging also significantly predicted all social threats such that higher levels were associated with higher perceived threat in general. Post-hoc evaluation determined this effect was only significantly different from zero for control (*p* = 0.0037; Figure 3D). There were no significant effects of mindset on self-reported social threats (*p* > 0.05; Table 2).

**Figure 3.**
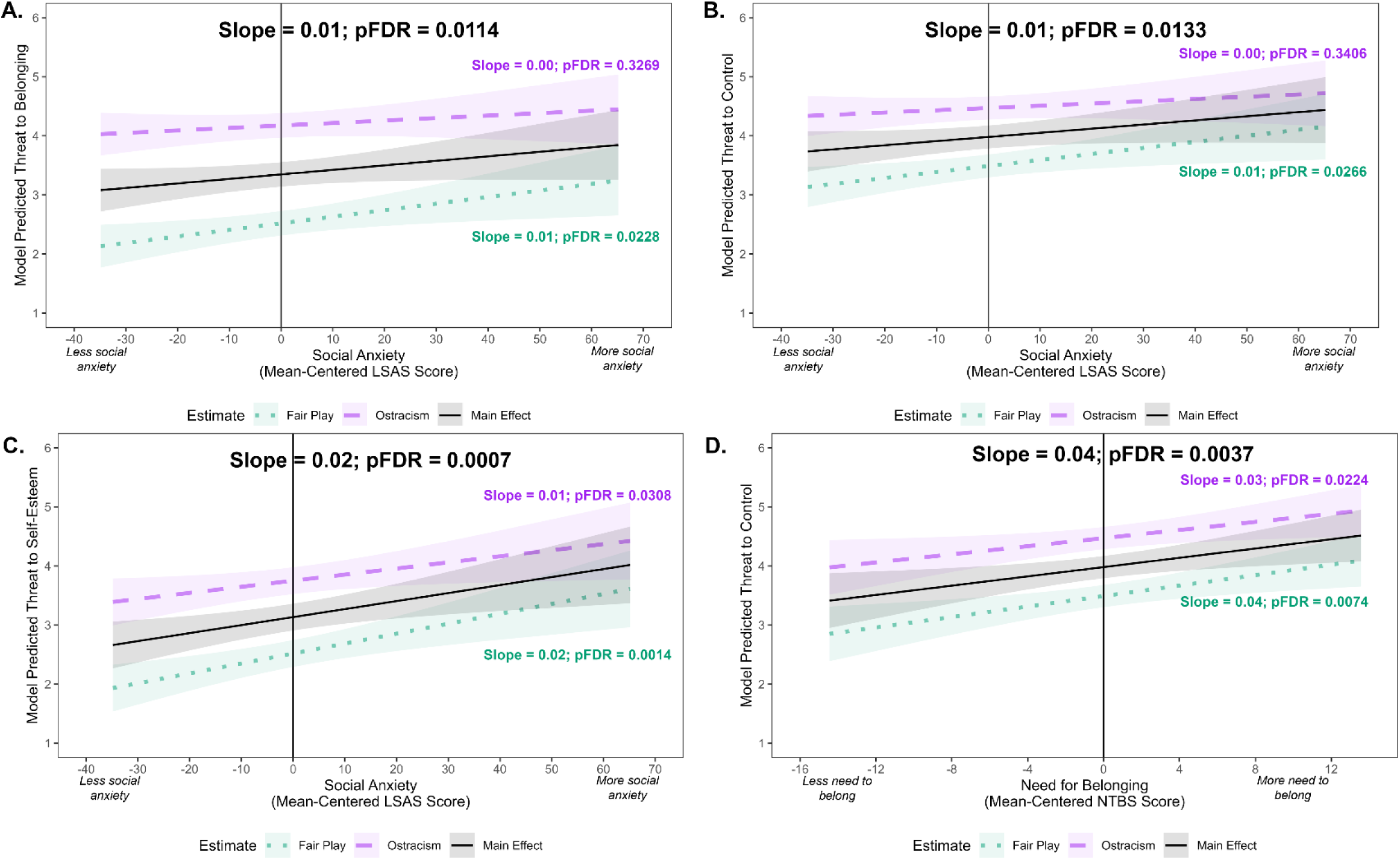
Individual difference effects illustrated by model-predicted self-reported sensitivity to social threats, as reported after completing each block. Across all panels slope estimates and FDR-corrected *p*-values are plotted. Turquoise dotted lines indicate predicted slopes for the fair play block and purple dashed lines indicate predicted slopes for the ostracism block. Black lines indicate predicted slopes for the main effect collapsed across both blocks. Horizontal lines indicate the mean score on each individual difference measure. **<u>Panel A-C</u>** represents the predicted relationship between social anxiety and social threats to belonging, control, and self-esteem respectively. **<u>Panel D</u>** represents the predicted relationship between need for belonging and social threats to control. LSAS = Liebowitz Social Anxiety Scale; NTBS = Need to Belong Scale

**Table 2.** Model estimates for block and individual difference effects on self-reported need-threat.

| Social Anxiety | Social Threat | Term | Fixed Effects |  |  | Interaction with Block |  |
| --- | --- | --- | --- | --- | --- | --- | --- |
|  |  |  | DF | F-value | p-value | F-value | p-value |
| Social Anxiety | Belonging | Intercept | 1, 51 | 1558.48 | <0.0001 |  |  |
|  |  | Block | 1, 51 | 203.19 | <0.0001 |  |  |
|  |  | LSAS | 1, 51 | 4.80 | 0.0330 | 2.09 | 0.1539 |
|  | Control | Intercept | 1, 51 | 2284.32 | <0.0001 |  |  |
|  |  | Block | 1, 51 | 100.10 | <0.0001 |  |  |
|  |  | LSAS | 1, 51 | 4.19 | 0.0459 | 2.49 | 0.1204 |
|  | Meaningful Existence | Intercept | 1, 51 | 879.49 | <0.0001 |  |  |
|  |  | Block | 1, 51 | 139.84 | <0.0001 |  |  |
|  |  | LSAS | 1, 51 | 4.51 | 0.0385 | 0.56 | 0.4571 |
|  | Self-Esteem | Intercept | 1, 51 | 1034.79 | <0.0001 |  |  |
|  |  | Block | 1, 51 | 116.64 | <0.0001 |  |  |
|  |  | LSAS | 1, 51 | 11.43 | 0.0014 | 1.90 | 0.1743 |
| Need for Belonging | Belonging | Intercept | 1, 51 | 1607.83 | <0.0001 |  |  |
|  |  | Block | 1, 51 | 195.82 | <0.0001 |  |  |
|  |  | NTBS | 1, 51 | 6.57 | 0.0134 | 0.17 | 0.6837 |
|  | Control | Intercept | 1, 51 | 2533.21 | <0.0001 |  |  |
|  |  | Block | 1, 51 | 96.20 | <0.0001 |  |  |
|  |  | NTBS | 1, 51 | 10.19 | 0.0024 | 0.41 | 0.5233 |
|  | Meaningful Existence | Intercept | 1, 51 | 886.55 | <0.0001 |  |  |
|  |  | Block | 1, 51 | 139.02 | <0.0001 |  |  |
|  |  | NTBS | 1, 51 | 4.96 | 0.0304 | 0.26 | 0.6142 |
|  | Self-Esteem | Intercept | 1, 51 | 936.99 | <0.0001 |  |  |
|  |  | Block | 1, 51 | 113.24 | <0.0001 |  |  |
|  |  | NTBS | 1, 51 | 5.53 | 0.0226 | 0.36 | 0.5532 |
| Personality Growth Mindset | Belonging | Intercept | 1, 51 | 1483.70 | <0.0001 |  |  |
|  |  | Block | 1, 51 | 195.59 | <0.0001 |  |  |
|  |  | ITP | 1, 51 | 2.12 | 0.1511 | 0.11 | 0.7427 |
|  | Control | Intercept | 1, 51 | 2192.50 | <0.0001 |  |  |
|  |  | Block | 1, 51 | 96.31 | <0.0001 |  |  |
|  |  | ITP | 1, 51 | 1.97 | 0.1667 | 0.46 | 0.4983 |
|  | Meaningful Existence | Intercept | 1, 51 | 829.75 | <0.0001 |  |  |
|  |  | Block | 1, 51 | 140.84 | <0.0001 |  |  |
|  |  | ITP | 1, 51 | 1.38 | 0.2461 | 0.92 | 0.3406 |
|  | Self-Esteem | Intercept | 1, 51 | 848.81 | <0.0001 |  |  |
|  |  | Block | 1, 51 | 113.33 | <0.0001 |  |  |
|  |  | ITP | 1, 51 | 0.20 | 0.6520 | 0.40 | 0.5294 |

**Note.** Individual models were run for each outcome measure and individual difference measure. LSAS = Liebowitz Social Anxiety Scale; NTBS = Need to Belong Scale; ITP = Implicit Theories of Personality

### 3.3. Differences in vlPFC activity between fair play and ostracism blocks

There was a significant effect of block on *z*-scored ΔHbO, *F*(1, 15655) = 7.83, *p* = 0.0051, such that the ostracism block elicited more vlPFC activity (EMM = -0.05, SE = 0.18) compared to the fair play block (EMM = -0.78, SE = 0.22; *p* = 0.0051). There was no significant effect of hemisphere, *F*(1, 52) = 0.01, *p* = 0.93 nor the interaction between hemisphere and block, *F*(1, 15655) = 0.76, *p* = 0.38, suggesting that the effect of block was widespread across both hemispheres (see Figure 4 and Supplemental Figure 1). Although separate models including self-reported social threats and individual difference measures indicated several initial significant main effects and interactions (see Supplemental Tables 4-6), post-hoc evaluation revealed that no relationships remained significant after FDR correction (*p* > 0.05).

**Figure 4.**
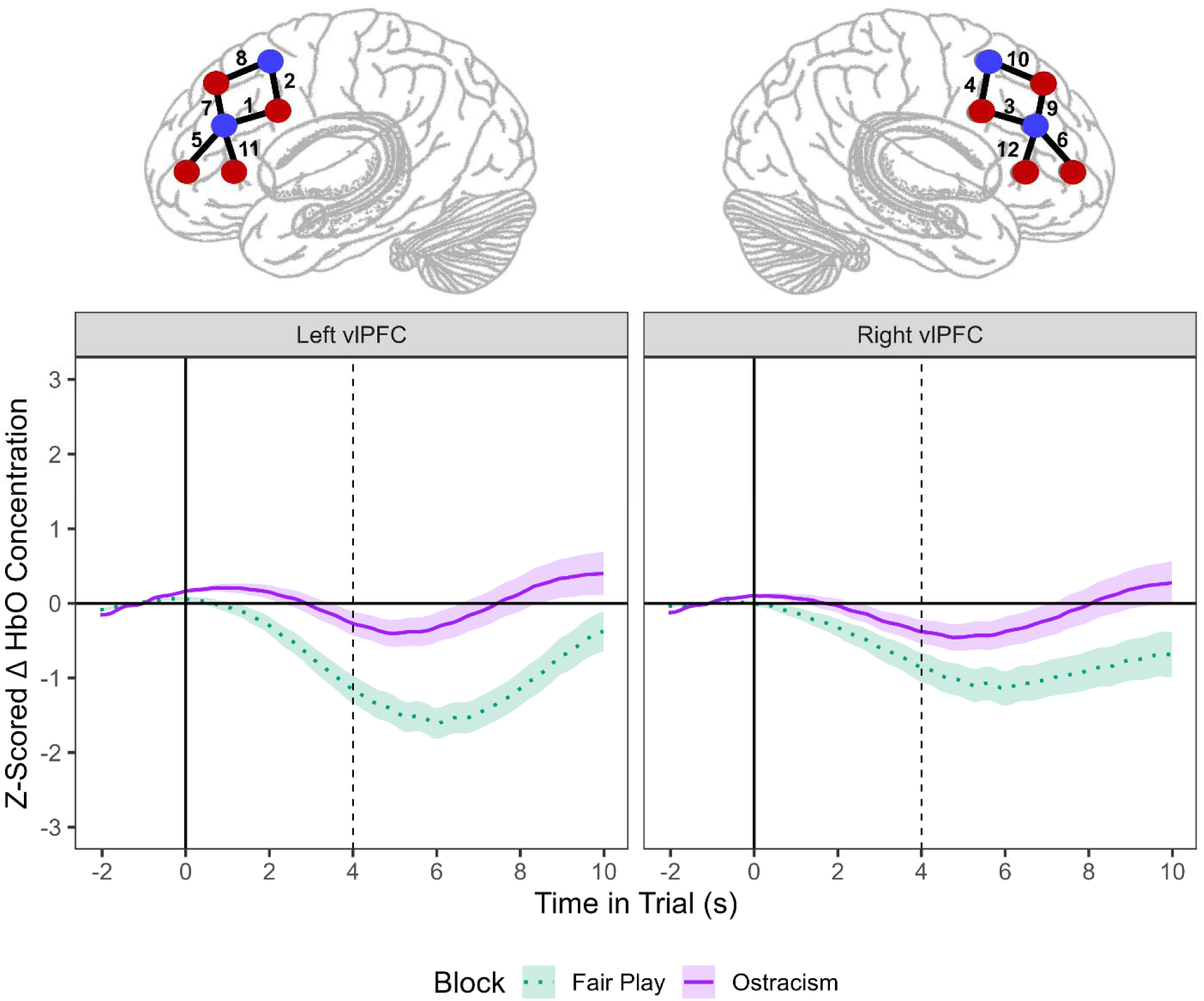
Block ΔHbO concentration changes over -2 to 10 s plotted across hemispheres. Across both panels, turquoise dotted lines represent *z*-scored ΔHbO across the fair play block and purple solid lines represent *z*-scored ΔHbO across the ostracism block. Dashed vertical lines indicate the end of a trial. **<u>Left panel</u>** represents data collapsed across channels located on the left hemisphere. **<u>Right panel</u>** represents data collapsed across channels located on the right hemisphere

### 3.4. vlPFC sensitization across inclusion and exclusion phases

Results from the model separating phase from block (i.e., fair play block inclusion phase, ostracism block inclusion phase, and ostracism block exclusion phase) indicated two significant two-way interactions between phase and trial number and trial number and hemisphere on *z*-scored ΔHbO (Table 3). Post-hoc evaluation revealed that trial number only had a significant effect during the ostracism exclusion phase such that vlPFC activity increased across trial number (estimate = 0.28; *p* < 0.0001). Additionally, this effect was slightly less positive for right vlPFC (estimate = 0.20) compared to left vlPFC (estimate = 0.37; *p* = 0.0044; Figure 5).

**Figure 5.**
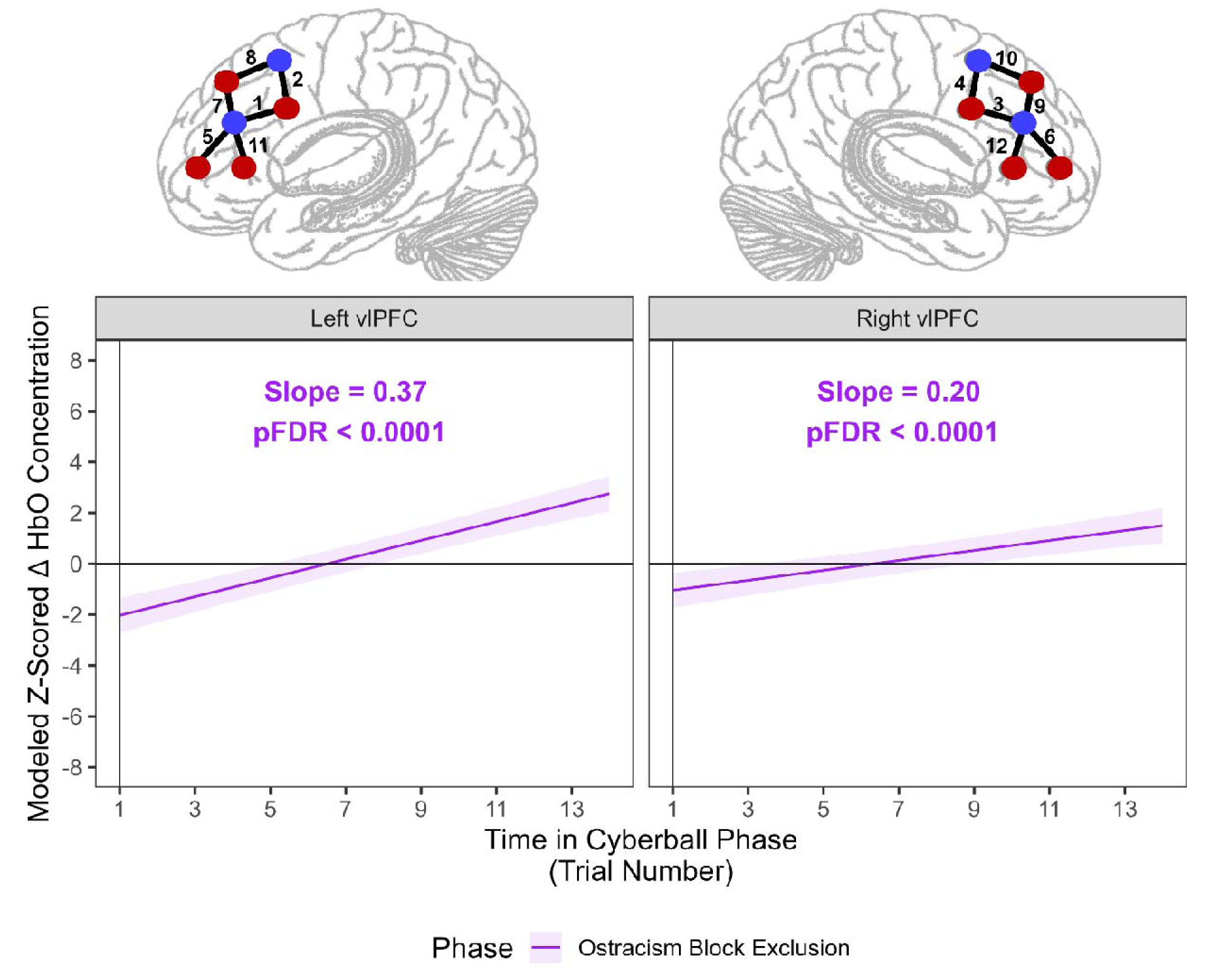
Effect of time on model-predicted *z*-scored ΔHbO concentration across phase and hemisphere. Across both panels, purple solid lines represent model-predicted linear ΔHbO *z*-scores across ostracism exclusion trials. Slope estimates and FDR-corrected p-values are plotted as well. **<u>Left panel</u>** represents model-predicted data from channels located on the left hemisphere. **<u>Right panel</u>** represents model-predicted data from channels located on the right hemisphere.

**Table 3.**
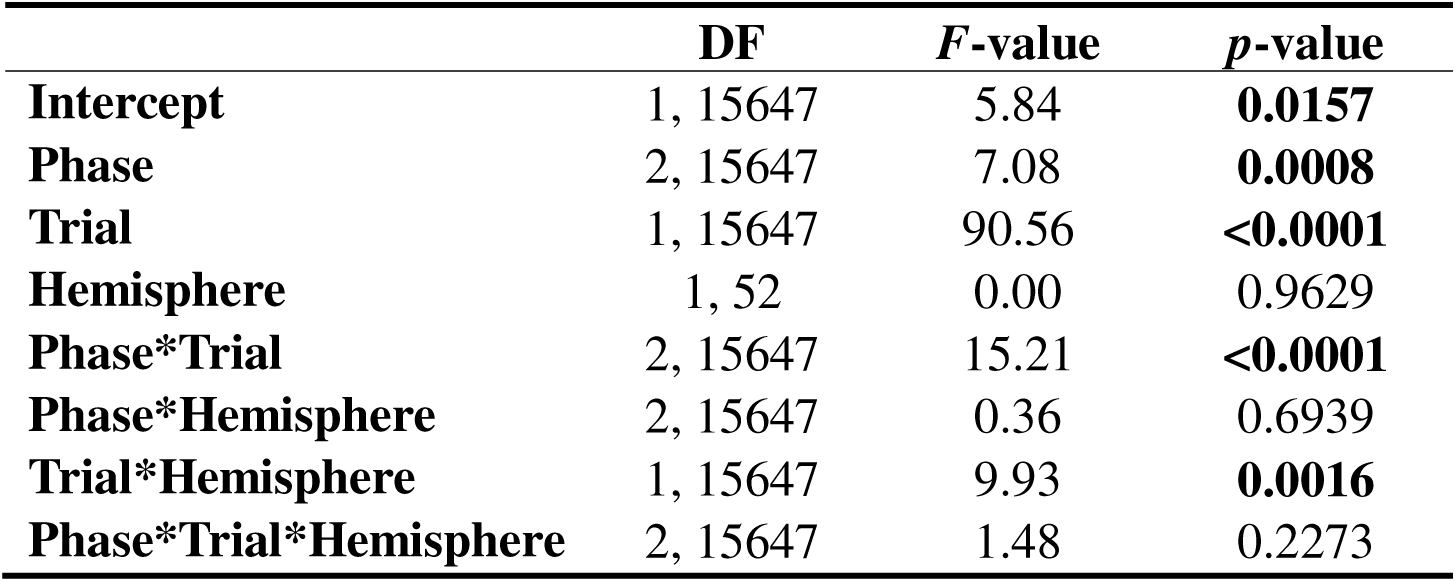
Model estimates for phase, hemisphere, and trial on ΔHbO *z*-scores.

|  | **DF** | ***F*-value** | ***p*-value** |
| --- | --- | --- | --- |
| **Intercept** | 1, 15647 | 5.84 | **0.0157** |
| **Phase** | 2, 15647 | 7.08 | **0.0008** |
| **Trial** | 1, 15647 | 90.56 | **<0.0001** |
| **Hemisphere** | 1, 52 | 0.00 | 0.9629 |
| **Phase\*Trial** | 2, 15647 | 15.21 | **<0.0001** |
| **Phase\*Hemisphere** | 2, 15647 | 0.36 | 0.6939 |
| **Trial\*Hemisphere** | 1, 15647 | 9.93 | **0.0016** |
| **Phase\*Trial\*Hemisphere** | 2, 15647 | 1.48 | 0.2273 |

### 3.5. The influence of individual differences on vlPFC sensitization to exclusion

For simplicity, we report only on significant interactions, although full model results can be found in Table 4. All individual difference variables were entered as continuous mean-centered predictors, though illustrations and post-hoc tests use representative levels (e.g., -1 SD below mean, mean, +1 SD above mean).

**Table 4.**
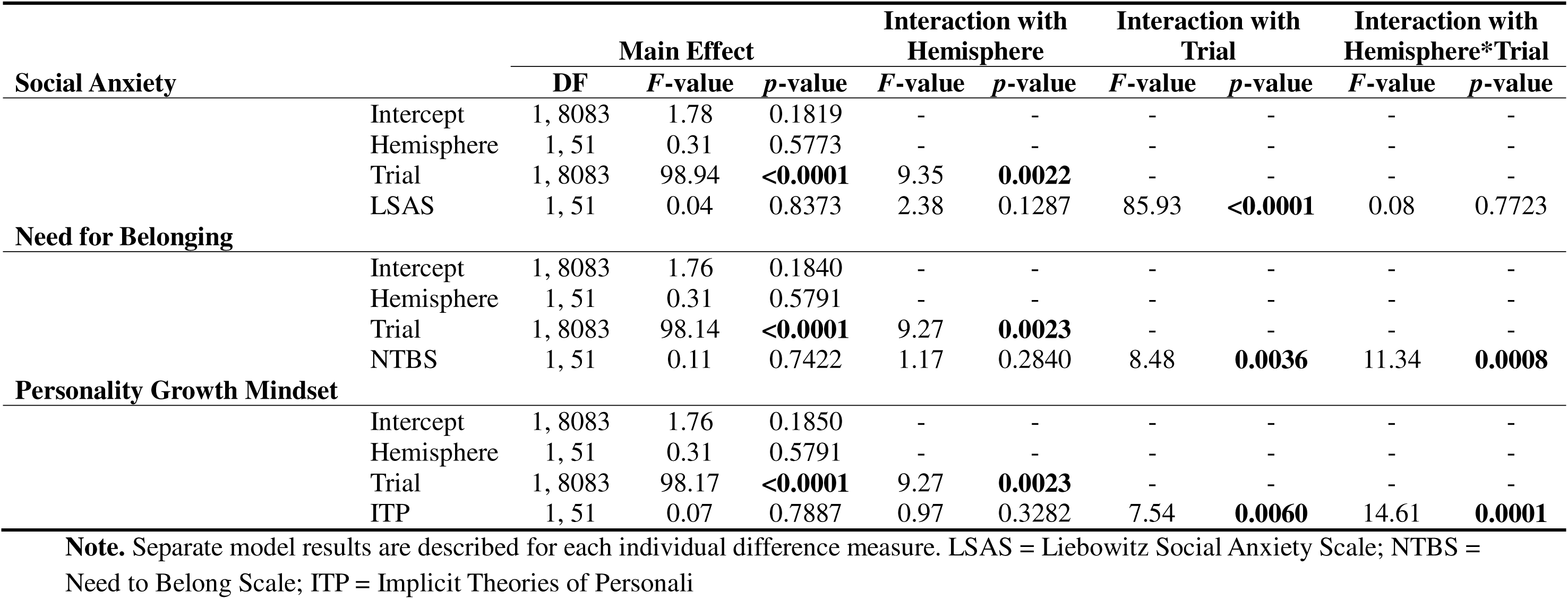
Model estimates for channel, trial, and individual difference measures on ΔHbO *z*-scores.

|  |  | Main Effect |  |  | Interaction with Hemisphere |  | Interaction with Trial |  | Interaction with Hemisphere*Trial |  |
| --- | --- | --- | --- | --- | --- | --- | --- | --- | --- | --- |
|  |  | DF | F-value | p-value | F-value | p-value | F-value | p-value | F-value | p-value |
| Social Anxiety | Intercept | 1, 8083 | 1.78 | 0.1819 | - | - | - | - | - | - |
|  | Hemisphere | 1, 51 | 0.31 | 0.5773 | - | - | - | - | - | - |
|  | Trial | 1, 8083 | 98.94 | <0.0001 | 9.35 | 0.0022 | - | - | - | - |
|  | LSAS | 1, 51 | 0.04 | 0.8373 | 2.38 | 0.1287 | 85.93 | <0.0001 | 0.08 | 0.7723 |
| Need for Belonging | Intercept | 1, 8083 | 1.76 | 0.1840 | - | - | - | - | - | - |
|  | Hemisphere | 1, 51 | 0.31 | 0.5791 | - | - | - | - | - | - |
|  | Trial | 1, 8083 | 98.14 | <0.0001 | 9.27 | 0.0023 | - | - | - | - |
|  | NTBS | 1, 51 | 0.11 | 0.7422 | 1.17 | 0.2840 | 8.48 | 0.0036 | 11.34 | 0.0008 |
| Personality Growth Mindset | Intercept | 1, 8083 | 1.76 | 0.1850 | - | - | - | - | - | - |
|  | Hemisphere | 1, 51 | 0.31 | 0.5791 | - | - | - | - | - | - |
|  | Trial | 1, 8083 | 98.17 | <0.0001 | 9.27 | 0.0023 | - | - | - | - |
|  | ITP | 1, 51 | 0.07 | 0.7887 | 0.97 | 0.3282 | 7.54 | 0.0060 | 14.61 | 0.0001 |

**Note.** Separate model results are described for each individual difference measure. LSAS = Liebowitz Social Anxiety Scale; NTBS = Need to Belong Scale; ITP = Implicit Theories of Personal

A significant two-way interaction between social anxiety and trial indicated that social anxiety influenced overall vlPFC sensitization (estimate = -0.01, *p* < 0.0001). Post-hoc analyses showed that individuals with severe social anxiety exhibited less neural sensitization to exclusion than those with no or mild social anxiety (*p* < 0.0001). Although individuals high in social anxiety began the exclusion phase with more positive vlPFC *z*-scored ΔHbO, they demonstrated less positive activity by the end of the phase (Figure 6A).

**Figure 6.**
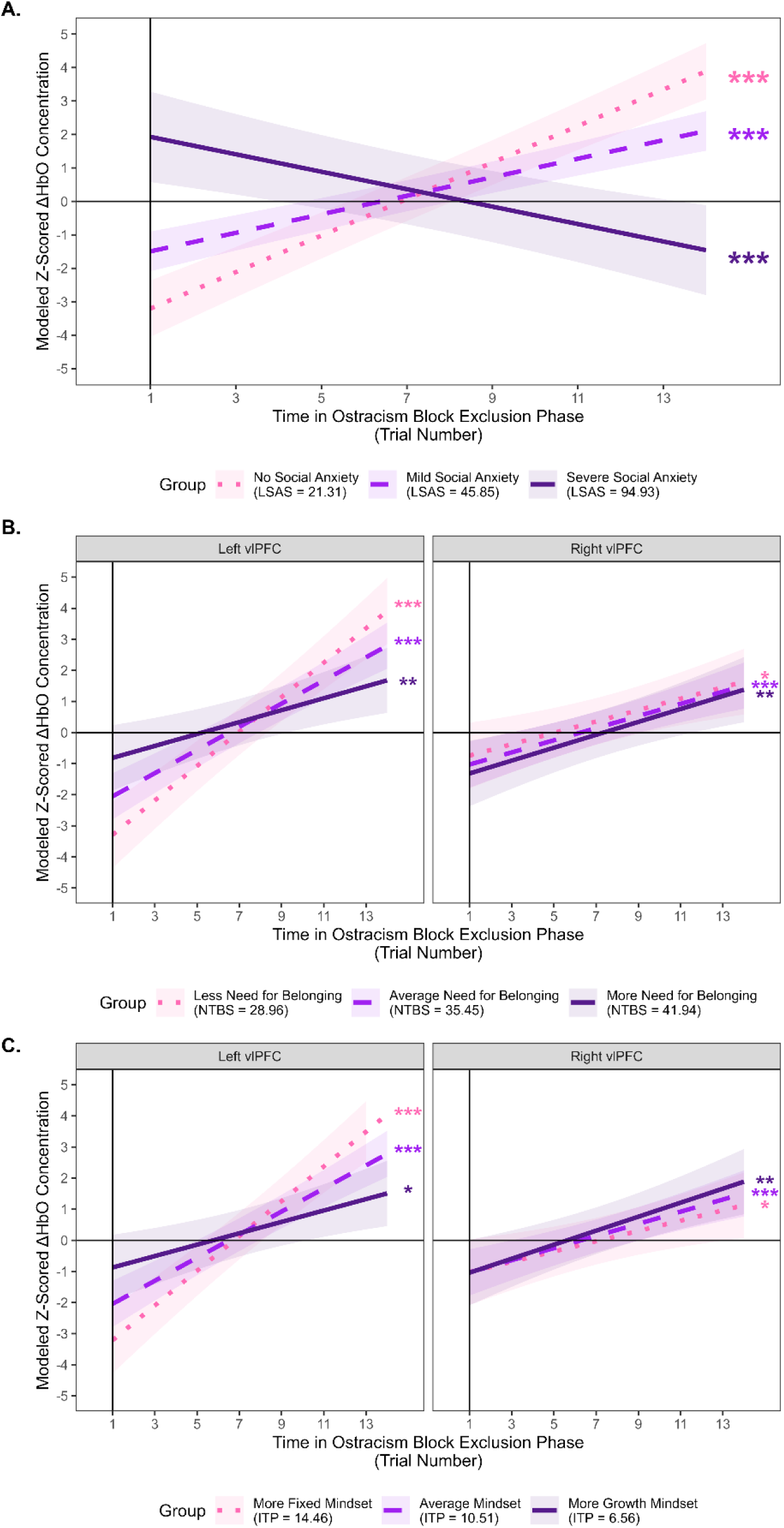
Plots of significant interactions between individual difference measures and trial on *z*-scored ΔHbO. Across all panels pink dotted lines indicate lower scores on the measure (e.g., -1 *SD* below mean values). Purple dashed lines indicate average scores on the measure (i.e., sample mean). Darker purple solid lines indicate higher scores on the measure (e.g., +1 *SD* above mean values). **<u>Panel A</u>** represents the linear effect of social anxiety and trial on model-predicted *z*-scored ΔHbO collapsed across both hemispheres. **<u>Panel B</u>** represents the linear effect of need for belonging and trial on model-predicted *z-*scored ΔHbO at left and right hemispheres. **<u>Panel C</u>** represents the linear effect of personality growth mindset and trial on model-predicted *z*-scored ΔHbO at left and right hemispheres. LSAS = Liebowitz Social Anxiety; NTBS = Need to Belong Scale; ITP = Implicit Theories of Personality; \**p* < 0.01; \*\**p* < 0.001; \*\*\**p* < 0.0001

A significant three-way interaction between need for belonging, trial, and hemisphere revealed hemispheric differences in how need for belonging influenced vlPFC sensitization (estimate = 0.03, *p* = 0.0008). Post-hoc tests demonstrated that right vlPFC sensitization increased over time regardless of need for belonging level (*p* = 0.95). In contrast, left vlPFC sensitization was strongest for those low in need for belonging compared to those at average or higher levels (*p* < 0.0001). Thus, although individuals with higher need for belonging began the exclusion phase with more positive left vlPFC *z*-scored ΔHbO, they showed less positive left vlPFC activity by the end of the phase (Figure 6B).

Finally, a significant three-way interaction between mindset, trial, and hemisphere indicated that mindset also differentially modulated hemispheric vlPFC sensitization (estimate = -0.05, *p* = 0.0001). Post-hoc analyses again showed increasing right vlPFC sensitization across all mindset levels (*p* = 0.57). In contrast, left vlPFC sensitization was highest for individuals with a more fixed mindset compared to those with more growth-oriented mindsets (*p* < 0.0001). In other words, individuals with greater growth mindset began the exclusion phase with more positive left vlPFC *z*-scored ΔHbO but ended with less positive activity than those with more fixed mindsets (Figure 6C).

## 4. Discussion

The present study confirmed that neural responses to ostracism increase as exclusion is experienced and that individual differences related to social anxiety, need for belonging, and mindset modulate behavioral and neural responses to ostracism. In line with classic Cyberball studies, we found the ostracism block predicted heightened self-reported social threats and vlPFC activity compared to the fair play block. Notably, increasing vlPFC sensitization emerged only during the ostracism block exclusion phase, with no comparable change in either inclusion phase. Although social anxiety and need for belonging modulated self-reported social threats, they did not predict overall vlPFC differences between the fair play and ostracism block. However, social anxiety, need for belonging, and mindset each predicted neural sensitization, such that individuals differing on these characteristics showed distinct patterns of vlPFC activity as exclusion was experienced.

### 4.1. Intensity of self-reported social threats is modulated by individual traits

Although all self-reported social threats were greater following the ostracism block compared to the fair play block, social anxiety and need for belonging modulated the intensity of social threats more generally. In other words, individuals who reported having more social anxiety and need for belonging reported increased social threats in both fair play and ostracism blocks. This result was unexpected given that research has demonstrated personality disorder traits and fear of social threat moderate self-reported distress for ostracism blocks, specifically (Riva et al., 2014; Wirth et al., 2010). Still, some research aligns with the present findings. For instance, Boyes and French (2009) found that high neuroticism participants reported less hedonic tone and self-esteem regardless of exclusion or inclusion. As such, it may be that whether individual differences influence exclusion-specific social threats may depend on the particular trait examined, though additional research is needed to clarify these distinctions.

### 4.2 Extending classic ostracism effects using single-trial analyses to uncover interpersonal differences in neural sensitization

Consistent with other classic neuroimaging work using the Cyberball paradigm (Eisenberger et al., 2003; Masten et al., 2009, 2011; Onoda et al., 2009, 2010), vlPFC activity across both hemispheres was greater for the ostracism block compared to the fair play block. Although no significant differences between right and left vlPFC may suggest that both right and left vlPFC are responsive to ostracism and may contribute to the ongoing debate about hemispheric differences in vlPFC activity related to emotion regulation (Cheng et al., 2022), these results should be interpreted cautiously. fNIRS offers better spatial resolution than methods such as electroencephalography (EEG), but its resolution remains lower than that of fMRI, and more channels are needed for more precise estimates of vlPFC activity (Chen et al., 2020). Considering that distinct hemispheric roles may reflect different mechanisms supporting emotion regulation during ostracism, clarifying the functions of the right and left vlPFC, alongside identifying effective regulation strategies, is crucial for understanding how the brain mitigates hypersensitivity to ostracism and downstream mental health consequences.

We found initial evidence for individual differences, social-threat, and the interactions between them predicting differences in vlPFC activity between the fair play and ostracism blocks. However, none of these relationships remained significant following corrections for multiple comparisons. While this was surprising considering neuroimaging research has found that increased self-reported social threats and individual differences predict vlPFC activity (Eisenberger et al., 2003; Gorka et al., 2018; Masten et al., 2009; Onoda et al., 2010), we kept blocks around two to three minutes like other fMRI work (Masten et al., 2011) which may not have been sufficient time to detect nuanced differences between block-level vlPFC activity using fNIRS. However, because of our single-trial fNIRS approach we were able to track vlPFC activity across three phases (i.e., fair play inclusion, ostracism inclusion, ostracism exclusion) and identified nuances in neural sensitization. Notably, vlPFC activity became increasingly positive only during the ostracism block exclusion phase. Given the vlPFC’s established role in emotion regulation (Ochsner et al., 2012) and in response to ostracism specifically (Eisenberger et al., 2003; Masten et al., 2009; Onoda et al., 2009), this result may suggest that as individuals become increasingly aware that they are being excluded they engage more resources to regulate emerging negative affect.

Critically, sensitization varied at the person-level, indicating that individual differences in social anxiety, need for belonging, and mindset shape emotion regulation following ostracism. Greater social anxiety was associated with less sensitization of vlPFC during the exclusion phase, suggesting that socially anxious individuals did not show increasing neural sensitivity as exclusion became more apparent. Although contrary to our hypotheses, prior research indicates that highly anxious individuals recover from social threats more slowly (Zadro et al., 2006). Additionally, individuals with social anxiety may also use different emotion regulation strategies compared to neurotypical controls (Dryman & Heimberg, 2018). As the vlPFC supports emotion regulation, reduced sensitization in more anxious participants may reflect weaker regulation or regulation recruiting non-vlPFC regions that may be associated with delayed recovery, though this was not directly tested here. Importantly, participants were not instructed to regulate their emotions, nor were emotion regulation strategies assessed. Anxious individuals typically recruit the vlPFC more when explicitly instructed to regulate negative affect (Campbell-Sills et al., 2011), raising the possibility that those higher in social anxiety were not regulating but instead ruminating—a hallmark of anxiety (Michl et al., 2013). Consistent with this, individuals with social anxiety disorder show reduced PFC activity in response to social threats (Goldin et al., 2009). Given that Cyberball exclusion is framed as real social evaluation, stable vlPFC activity in socially anxious participants may reflect recognition of real social threats. Finally, social anxiety scores for the current sample mostly fell in the “no social anxiety” or “mild social anxiety” groups. Considering ostracism and social anxiety may mutually reinforce one another (Albath et al., 2025; Büttner & Greifeneder, 2024; Reinhard et al., 2020; Rudert et al., 2021), future research should continue examining how social anxiety traits relate to vlPFC sensitization over the course of ostracism experiences, especially in clinically elevated populations.

Need for belonging also modulated vlPFC sensitization during the exclusion phase, however the pattern was different across hemispheres. Although right vlPFC activity increased for all levels of need for belonging across the exclusion phase, individuals with more need for belonging demonstrated less sensitization in left vlPFC compared to individuals with less need for belonging. Given that need for belonging has been associated with negative social emotions, neuroticism, and rejection sensitivity (Leary et al., 2013), it is reasonable that individual differences in this trait shaped vlPFC activity as exclusion progressed. Although hemisphere was not a significant predictor in the preceding models, it appears to contribute meaningfully here. Indeed, differentiated hemispheric activity may align with research suggesting functional hemispheric differences in emotion regulation (Cheng et al., 2022). For instance, disrupting left vlPFC activity reduces semantic processing of emotions, whereas disrupting right vlPFC reduces inhibitory downregulation of negative affect (Cheng et al., 2022). Thus, in the present study, individuals with greater need for belonging may rely less on semantic related strategies and more on inhibitory regulation strategies during ostracism. Notably, even though left vlPFC change was attenuated, slope estimates remained positive, indicating that overall vlPFC activity was increasing with higher need for belonging. Continued examination of hemispheric vlPFC dynamics during ostracism will be important for clarifying these relationships.

Personality growth mindset also modulated vlPFC sensitization, with effects differing by hemisphere. Right vlPFC activity increased across the exclusion phase for all participants, whereas individuals with stronger growth mindset showed less left vlPFC sensitization than those with a fixed mindset. This pattern further supports that hemispheric vlPFC functions may depend on person-level traits. Personality growth mindset has been linked to reduced retaliatory desires following peer victimization (Yeager et al., 2011) and to mediating associations among self-esteem, ostracism, and aggression (Li et al., 2019). Since right vlPFC activity is associated with inhibitory emotion regulation (Cheng et al., 2022) and poor inhibitory control is associated with higher trait aggression levels (Pawliczek et al., 2013; Romero-López et al., 2021), the pattern identified here may indicate that growth-minded individuals engage more inhibitory regulation during ostracism. Although we did not measure aggression or retaliatory tendencies, future work should aim to better clarify this relationship. Neuroimaging research on growth mindset is limited (Ng, 2018), but existing studies implicate brain regions involved in conflict monitoring that are also related to ostracism processing (e.g., ACC; dlPFC; (Myers et al., 2016). Growth mindset interventions have also proven to improve academic outcomes (Blackwell et al., 2007), increase academic engagement (Martin et al., 2019), reduce retaliatory desires following peer victimization (Yeager et al., 2011), and increase vlPFC activity during response inhibition (Blanchette Sarrasin et al., 2025). Further research would benefit from elucidating how growth mindset relates to neural mechanisms of conflict monitoring and emotion regulation, and whether interventions might buffer against heightened sensitivity to ostracism. Finally, because social anxiety, need for belonging, and mindset were not correlated in our sample, we examined their effects separately; however, their theoretical overlap and influence on ostracism suggests future work should assess their joint influence on vlPFC sensitization during social exclusion.

## 5. Conclusions

Using a single-trial approach, the present study demonstrates that vlPFC sensitivity progressively increases with repeated experiences of ostracism, revealing previously uncharacterized temporal dynamics in neural responses to social exclusion. Notably, these dynamics were systematically predicted by person-level traits, highlighting meaningful individual differences in neural sensitivity to ostracism. Given evidence that heightened sensitivity to ostracism is associated with increased risk for adverse mental health outcomes, the current findings provide a critical mechanistic link between individual traits and neural sensitization processes. By identifying differentiated neural sensitization patterns associated with social exclusion, this work offers a foundation for understanding how maladaptive responses to ostracism may emerge and persist over time.

## Acknowledgements

We would like to thank our participants for their willingness to participate in our study. We also would like to acknowledge [anonymized for review] for their assistance in creating stimuli for and piloting the task and reviewing earlier drafts of this manuscript. The research was partially supported by The Center for the Study of Ethical Development at the University of Alabama.

## Notes

### Competing Interest Statement

The authors have declared no competing interest.

## References

Albath, E. A., Bogatyreva, N., Büttner, C. M., & Greifeneder, R. (2025). The role of depression and anxiety in experiencing and inflicting ostracism: A cross-national perspective. Journal of Affective Disorders, 380, 696–703. 10.1016/j.jad.2025.03.190

Bates, D., Mächler, M., Bolker, B., & Walker, S. (2015). Fitting Linear Mixed-Effects Models Using lme4. Journal of Statistical Software, 67(1). 10.18637/jss.v067.i01

Baumeister, R. F., & Leary, M. R. (1995). The Need to Belong: Desire for Interpersonal Attachments as a Fundamental Human Motivation. Psychollogical Bulletin, 117(3), 497–529.

Benjamini, Y., & Hochberg, Y. (1995). Controlling the false discovery rate: A practical and powerful approach to multiple testing. Journal of the Royal Statistical Society Series B (Methodological), 57(1), 289–300.

Blackwell, L. S., Trzesniewski, K. H., & Dweck, C. S. (2007). Implicit Theories of Intelligence Predict Achievement Across an Adolescent Transition: A Longitudinal Study and an Intervention. Child Development, 78(1), 246–263. 10.1111/j.1467-8624.2007.00995.x

Blanchette Sarrasin, J., Riopel, M., Allaire-Duquette, G., McMullin, S., Bélanger, É., Brault Foisy, L. M., & Masson, S. (2025). Effects of teaching neuroplasticity on motivation, inhibitory control and task performance, and the role of mindset theory. Trends in Neuroscience and Education, 40. 10.1016/j.tine.2025.100257

Bolling, D. Z., Pitskel, N. B., Deen, B., Crowley, M. J., McPartland, J. C., Mayes, L. C., & Pelphrey, K. A. (2011). Dissociable brain mechanisms for processing social exclusion and rule violation. NeuroImage, 54(3), 2462–2471. 10.1016/j.neuroimage.2010.10.049

Boyes, M. E., & French, D. J. (2009). Having a Cyberball: Using a ball-throwing game as an experimental social stressor to examine the relationship between neuroticism and coping. Personality and Individual Differences, 47(5), 396–401. 10.1016/j.paid.2009.04.005

Büttner, C. M., & Greifeneder, R. (2024). Everyday ostracism experiences of depressed individuals: Uncovering the role of attributions using experience sampling. Journal of Affective Disorders Reports, 17. 10.1016/j.jadr.2024.100804

Cacioppo, S., Frum, C., Asp, E., Weiss, R. M., Lewis, J. W., & Cacioppo, J. T. (2013). A quantitative meta-analysis of functional imaging studies of social rejection. Scientific Reports, 3. 10.1038/srep02027

Campbell-Sills, L., Simmons, A. N., Lovero, K. L., Rochlin, A. A., Paulus, M. P., & Stein, M. B. (2011). Functioning of neural systems supporting emotion regulation in anxiety-prone individuals. NeuroImage, 54(1), 689–696. 10.1016/j.neuroimage.2010.07.041

Chen, W. L., Wagner, J., Heugel, N., Sugar, J., Lee, Y. W., Conant, L., Malloy, M., Heffernan, J., Quirk, B., Zinos, A., Beardsley, S. A., Prost, R., & Whelan, H. T. (2020). Functional Near-Infrared Spectroscopy and Its Clinical Application in the Field of Neuroscience: Advances and Future Directions. In Frontiers in Neuroscience (Vol. 14). Frontiers Media S.A. 10.3389/fnins.2020.00724

Cheng, S., Qiu, X., Li, S., Mo, L., Xu, F., & Zhang, D. (2022). Different Roles of the Left and Right Ventrolateral Prefrontal Cortex in Cognitive Reappraisal: An Online Transcranial Magnetic Stimulation Study. Frontiers in Human Neuroscience, 16. 10.3389/fnhum.2022.928077

Dou, H., Lei, Y., Cheng, X., Wang, J., & Leppänen, P. H. T. (2020). Social exclusion influences conditioned fear acquisition and generalization: A mediating effect from the medial prefrontal cortex. NeuroImage, 218. 10.1016/j.neuroimage.2020.116735

Dryman, M. T., & Heimberg, R. G. (2018). Emotion regulation in social anxiety and depression: a systematic review of expressive suppression and cognitive reappraisal. In Clinical Psychology Review (Vol. 65, pp. 17–42). Elsevier Inc. 10.1016/j.cpr.2018.07.004

Eisenberger, N. I., Lieberman, M. D., & Williams, K. D. (2003). Does rejection hurt? An fMRI study of social exclusion. Science, 302, 290–292.

Goldin, P. R., Manber, T., Hakimi, S., Canli, T., & Gross, J. J. (2009). Neural Bases of Social Anxiety Disorder. Archives of General Psychiatry, 66(2), 170. 10.1001/archgenpsychiatry.2008.525

Gorka, S. M., Phan, K. L., Hosseini, B., Chen, E. Y., & McCloskey, M. S. (2018). Ventrolateral Prefrontal Cortex Activation During Social Exclusion Mediates the Relation Between Intolerance of Uncertainty and Trait Aggression. Clinical Psychological Science, 6(6), 810–821. 10.1177/2167702618776947

Hales, A. H., Ren, D., & Williams, K. D. (2016). Protect, correct, and eject: Ostracism as a social influence tool. In S. J. Harkins, J. M. Burger, & K. D. Williams (Eds.), The Oxford Handbook of Social Influence (pp. 205–218). Oxford University Press.

Hartgerink, C. H. J., Van Beest, I., Wicherts, J. M., & Williams, K. D. (2015). The ordinal effects of ostracism: A meta-analysis of 120 cyberball studies. PLoS ONE, 10(5). 10.1371/journal.pone.0127002

Homan, R. W., Herman, J., & Purdy, P. (1987). Cerebral location of international 10–20 system electrode placement. Electroencephalography and Clinical Neurophysiology, 66(4), 376–382. 10.1016/0013-4694(87)90206-9

Huppert, T. J., Diamond, S. G., Franceschini, M. A., & Boas, D. A. (2009). HomER: a review of time-series analysis methods for near-infrared spectroscopy of the brain. Applied Optics, 48(10), D280. 10.1364/AO.48.00D280

Jahani, S., Setarehdan, S. K., Boas, D. A., & Yücel, M. A. (2018). Motion artifact detection and correction in functional near-infrared spectroscopy: a new hybrid method based on spline interpolation method and Savitzky–Golay filtering. Neurophotonics, 5(01), 1. 10.1117/1.NPh.5.1.015003

Jiao, Z., Song, J., Yang, X., Chen, Y., & Han, G. (2024). Social pain sharing boosts interpersonal brain synchronization in female cooperation. Acta Psychologica, 243. 10.1016/j.actpsy.2024.104138

Klein, F., Lührs, M., Benitez-Andonegui, A., Roehn, P., & Kranczioch, C. (2022). Performance comparison of systemic activity correction in functional near-infrared spectroscopy for methods with and without short distance channels. Neurophotonics, 10(01). 10.1117/1.NPh.10.1.013503

Leary, M. R., Kelly, K. M., Cottrell, C. A., & Schreindorfer, L. S. (2013). Construct validity of the need to belong scale: Mapping the nomological network. Journal of Personality Assessment, 95(6), 610–624. 10.1080/00223891.2013.819511

Li, S., Xie, H., Zheng, Z., Chen, W., Xu, F., Hu, X., & Zhang, D. (2022). The causal role of the bilateral ventrolateral prefrontal cortices on emotion regulation of social feedback. Human Brain Mapping, 43(9), 2898–2910. 10.1002/hbm.25824

Li, S., Zhao, F., & Yu, G. (2019). Ostracism and aggression among adolescents: Implicit theories of personality moderated the mediating effect of self-esteem. Children and Youth Services Review, 100, 105–111. 10.1016/j.childyouth.2019.02.043

Lian, T., Jiao, Z., Juan, S., & Zhang, P. (2025). Interpersonal brain synchronization in social pain contexts: an fNIRS-based exploration of empathy. Social Cognitive and Affective Neuroscience, 20(1). 10.1093/scan/nsaf003

Lieberman, M. D. (2007). Social cognitive neuroscience: A review of core processes. In Annual Review of Psychology (Vol. 58, pp. 259–289). 10.1146/annurev.psych.58.110405.085654

Liebowitz, M. R. (1987). Social Phobia. Modern Problems in Pharmacopsychiatry, 22, 141–173. 10.1159/000414022

Martin, A. J., Collie, R. J., Durksen, T. L., Burns, E. C., Bostwick, K. C. P., & Tarbetsky, A. L. (2019). Growth goals and growth mindset from a methodological-synergistic perspective: lessons learned from a quantitative correlational research program. International Journal of Research & Method in Education, 42(2), 204–219. 10.1080/1743727X.2018.1481938

Masten, C. L., Colich, N. L., Rudie, J. D., Bookheimer, S. Y., Eisenberger, N. I., & Dapretto, M. (2011). An fMRI investigation of responses to peer rejection in adolescents with autism spectrum disorders. Developmental Cognitive Neuroscience, 1(3), 260–270. 10.1016/j.dcn.2011.01.004

Masten, C. L., Eisenberger, N. I., Borofsky, L. A., Pfeifer, J. H., McNealy, K., Mazziotta, J. C., & Dapretto, M. (2009). Neural correlates of social exclusion during adolescence: Understanding the distress of peer rejection. Social Cognitive and Affective Neuroscience, 4(2), 143–157. 10.1093/scan/nsp007

Michl, L. C., McLaughlin, K. A., Shepherd, K., & Nolen-Hoeksema, S. (2013). Rumination as a mechanism linking stressful life events to symptoms of depression and anxiety: Longitudinal evidence in early adolescents and adults. Journal of Abnormal Psychology, 122(2), 339–352. 10.1037/a0031994

Myers, C. A., Wang, C., Black, J. M., Bugescu, N., & Hoeft, F. (2016). The matter of motivation: Striatal resting-state connectivity is dissociable between grit and growth mindset. Social Cognitive and Affective Neuroscience, 11(10), 1521–1527. 10.1093/scan/nsw065

Ng, B. (2018). The Neuroscience of Growth Mindset and Intrinsic Motivation. Brain Sciences, 8(2), 20. 10.3390/brainsci8020020

Nishiyama, Y., Okamoto, Y., Kunisato, Y., Okada, G., Yoshimura, S., Kanai, Y., Yamamura, T., Yoshino, A., Jinnin, R., Takagaki, K., Onoda, K., & Yamawaki, S. (2015). fMRI study of social anxiety during social ostracism with and without emotional support. PLoS ONE, 10(5). 10.1371/journal.pone.0127426

Ochsner, K. N., Silvers, J. A., & Buhle, J. T. (2012). Functional imaging studies of emotion regulation: a synthetic review and evolving model of the cognitive control of emotion. In Annals of the New York Academy of Sciences (Vol. 1251, pp. E1–E24). 10.1111/j.1749-6632.2012.06751.x

Onoda, K., Okamoto, Y., Nakashima, K., Nittono, H., Ura, M., & Yamawaki, S. (2009). Decreased ventral anterior cingulate cortex activity is associated with reduced social pain during emotional support. Social Neuroscience, 4(5), 443–454. 10.1080/17470910902955884

Onoda, K., Okamoto, Y., Nakashima, K., Nittono, H., Yoshimura, S., Yamawaki, S., Yamaguchi, S., & Ura, M. (2010). Does low self-esteem enhance social pain? The relationship between trait self-esteem and anterior cingulate cortex activation induced by ostracism. Social Cognitive and Affective Neuroscience, 5(4), 385–391. 10.1093/scan/nsq002

O’Reilly, R. C. (2010). The What and How of prefrontal cortical organization. Trends in Neurosciences, 33(8), 355–361. 10.1016/j.tins.2010.05.002

Pawliczek, C. M., Derntl, B., Kellermann, T., Kohn, N., Gur, R. C., & Habel, U. (2013). Inhibitory control and trait aggression: Neural and behavioral insights using the emotional stop signal task. NeuroImage, 79, 264–274. 10.1016/j.neuroimage.2013.04.104

Reinhard, M. A., Dewald-Kaufmann, J., Wüstenberg, T., Musil, R., Barton, B. B., Jobst, A., & Padberg, F. (2020). The vicious circle of social exclusion and psychopathology: a systematic review of experimental ostracism research in psychiatric disorders. European Archives of Psychiatry and Clinical Neuroscience, 270(5), 521–532. 10.1007/s00406-019-01074-1

Riva, P., Marinucci, M., Telari, A., & Pancani, L. (2025). Updating the temporal need-threat model of ostracism: challenges and future directions. Journal of Social Psychology. 10.1080/00224545.2025.2572648

Riva, P., Williams, K. D., & Gallucci, M. (2014). The relationship between fear of social and physical threat and its effect on social distress and physical pain perception. Pain, 155(3), 485–493. 10.1016/j.pain.2013.11.006

Romero-López, M., Pichardo, M. C., Justicia-Arráez, A., & Bembibre-Serrano, J. (2021). Reducing Aggression by Developing Emotional and Inhibitory Control. International Journal of Environmental Research and Public Health, 18(10), 5263. 10.3390/ijerph18105263

Rudert, S. C., Janke, S., & Greifeneder, R. (2021). Ostracism breeds depression: Longitudinal associations between ostracism and depression over a three-year-period. Journal of Affective Disorders Reports, 4. 10.1016/j.jadr.2021.100118

Smith, A., & Williams, K. D. (2004). R u there? Ostracism by cell phone text messages. Group Dynamics, 8(4), 291–301. 10.1037/1089-2699.8.4.291

Song, J., Lian, T., Zhang, Y., Cao, M., & Jiao, Z. (2024). Social exclusion: differences in neural mechanisms underlying direct versus vicarious experience. Frontiers in Psychology, 15. 10.3389/fpsyg.2024.1368214

Stokes, M., & Spaak, E. (2016). The importance of single-trial analyses in cognitive neuroscience. Trends in Cognitive Sciences, 20(7), 483–486. 10.1016/j.tics.2016.04.005

Su, W. C., Dashtestani, H., Miguel, H. O., Condy, E., Buckley, A., Park, S., Perreault, J. B., Nguyen, T., Zeytinoglu, S., Millerhagen, J., Fox, N., & Gandjbakhche, A. (2023). Simultaneous multimodal fNIRS-EEG recordings reveal new insights in neural activity during motor execution, observation, and imagery. Scientific Reports, 13(1). 10.1038/s41598-023-31609-5

van Noordt, S. J. R., White, L. O., Wu, J., Mayes, L. C., & Crowley, M. J. (2015). Social exclusion modulates event-related frontal theta and tracks ostracism distress in children. NeuroImage, 118, 248–255. 10.1016/j.neuroimage.2015.05.085

von Lühmann, A., Ortega-Martinez, A., Boas, D. A., & Yücel, M. A. (2020). Using the General Linear Model to Improve Performance in fNIRS Single Trial Analysis and Classification: A Perspective. Frontiers in Human Neuroscience, 14. 10.3389/fnhum.2020.00030

Williams, K. D. (2009). Ostracism: A Temporal Need-Threat Model. In Advances in Experimental Social Psychology (Vol. 41, pp. 275–314). 10.1016/S0065-2601(08)00406-1

Williams, K. D., Cheung, C. K. T., & Choi, W. (2000). Cyberostracism: Effects of being ignored over the Internet. Journal of Personality and Social Psychology, 79(5), 748–762. 10.1037/0022-3514.79.5.748

Williams, K. D., & Sommer, K. L. (1997). Social ostracism by coworkers: Does rejection lead to loafing or compensation? Personality and Social Psychology Bulletin, 23(7), 693–706. 10.1177/0146167297237003

Wirth, J. H., Lynam, D. R., & Williams, K. D. (2010). When social pain is not automatic: Personality disorder traits buffer ostracism’s immediate negative impact. Journal of Research in Personality, 44(3), 397–401. 10.1016/j.jrp.2010.03.001

Yeager, D. S., & Dweck, C. S. (2012). Mindsets That Promote Resilience: When Students Believe That Personal Characteristics Can Be Developed. In Educational Psychologist (Vol. 47, Issue 4, pp. 302–314). 10.1080/00461520.2012.722805

Yeager, D. S., & Miu, A. (2011). Implicit theories of personality predict motivation to use prosocial coping strategies after bullying in high school. In E. Frydenberg & G. Reevy (Eds.), Personality, stress, and coping: Implications for education (pp. 47–62). Information Age.

Yeager, D. S., Trzesniewski, K. H., Tirri, K., Nokelainen, P., & Dweck, C. S. (2011). Adolescents’ Implicit Theories Predict Desire for Vengeance After Peer Conflicts: Correlational and Experimental Evidence. Developmental Psychology, 47(4), 1090–1107. 10.1037/a0023769

Yücel, M. A., Lühmann, A. v., Scholkmann, F., Gervain, J., Dan, I., Ayaz, H., Boas, D., Cooper, R. J., Culver, J., Elwell, C. E., Eggebrecht, A., Franceschini, M. A., Grova, C., Homae, F., Lesage, F., Obrig, H., Tachtsidis, I., Tak, S., Tong, Y., … Wolf, M. (2021). Best practices for fNIRS publications. Neurophotonics, 8(01). 10.1117/1.nph.8.1.012101

Zadro, L., Boland, C., & Richardson, R. (2006). How long does it last? The persistence of the effects of ostracism in the socially anxious. Journal of Experimental Social Psychology, 42(5), 692–697. 10.1016/j.jesp.2005.10.007

Zimeo Morais, G. A., Balardin, J. B., & Sato, J. R. (2018). FNIRS Optodes’ Location Decider (fOLD): A toolbox for probe arrangement guided by brain regions-of-interest. Scientific Reports, 8(1). 10.1038/s41598-018-21716-z

