## Supplementary Materials for "Individual traits shape hemispheric vlPFC sensitization to Cyberball exclusion: Evidence from single-trial fNIRS analyses"

**Supplemental Materials**

**1.1 R syntax for linear mixed-effects models predicting social threat responses measured by the Need Threat Survey (NTS) across block and individual difference measures.** Individual difference mean-centered scores (social anxiety [LSAS], need for belonging [NTBS], and mindset [ITP] are entered in separate models for each of the social threat variables (belonging, control, meaningful existence, self-esteem):

*lme(Social Threat Variable~Block*Individual Difference Mean-Centered Score, random =~1|id, data = df, na.action = na.exclude, method = “REML”, control = lmeControl(opt = “optim”)).*

**1.2 R syntax for linear mixed-effects models predicting z-scored changes in HbO by block, hemisphere, NTS responses, and individual differences.**

Model 1: *lme(HbO Z-Score~Block*Hemisphere*Mean-Centered Threat to Belonging*Mean-Centered Social Anxiety + Block*Hemisphere*Mean-Centered Threat to Control*Mean-Centered Social Anxiety+ Block*Hemisphere*Mean-Centered Threat to Meaningful Existence*Mean-Centered Social Anxiety+Block*Hemisphere*Mean-Centered Threat to Self-Esteem*Mean-Centered Social Anxiety, random =~1+Block|ID/Hemisphere, data = df, na.action = na.exclude, method = “REML”, control = list(opt = "optim"))*

Model 2: *lme(HbO Z-Score~Block*Hemisphere*Mean-Centered Threat to Belonging*Mean-Centered Need for Belonging + Block*Hemisphere*Mean-Centered Threat to Control*Mean-Centered Need for Belonging+ Block*Hemisphere*Mean-Centered Threat to Meaningful Existence*Mean-Centered Need for Belonging+Block*Hemisphere*Mean-Centered Threat to Self-Esteem*Mean-Centered Need for Belonging, random =~1+Block|ID/Hemisphere, data = df, na.action = na.exclude, method = “REML”, control = list(opt = "optim"))*

Model 3: *lme(HbO Z-Score~Block*Hemisphere*Mean-Centered Threat to Belonging*Mean-Centered Mindset + Block*Hemisphere*Mean-Centered Threat to Control*Mean-Centered Mindset+ Block*Hemisphere*Mean-Centered Threat to Meaningful Existence*Mean-Centered Mindset+Block*Hemisphere*Mean-Centered Threat to Self-Esteem*Mean-Centered Mindset, random =~1+Block|ID/Hemisphere, data = df, na.action = na.exclude, method = “REML”, control = list(opt = "optim"))*

**1.3 R syntax for linear mixed-effects model predicting z-scored changes in HbO within exclusion trials by hemisphere, trial, and individual difference measures.**

Model 1: *lme(Exclusion Phase HbO Z-Score~Trial Number*Hemisphere*Mean-Centered Social Anxiety, random =~1|ID/Hemisphere, data = df, na.action = na.exclude, method = “REML”, control = list(opt = "optim"))*

Model 2: *lme(Exclusion Phase HbO Z-Score~Trial Number*Hemisphere*Mean-Centered Need for Belonging, random =~1|ID/Hemisphere, data = df, na.action = na.exclude, method = “REML”, control = list(opt = "optim"))*

Model 3: *lme(Exclusion Phase HbO Z-Score~Trial Number*Hemisphere*Mean-Centered Mindset, random =~1|ID/Hemisphere, data = df, na.action = na.exclude, method = “REML”, control = list(opt = "optim"))*

**Supplemental Table 1. Model results from sensitivity analyses comparing inclusion trial lengths across blocks.**

| **Term** | **DF** | ***F*-value** | ***p*-value** |
| --- | --- | --- | --- |
| Intercept | 1, 17938 | 5.36 | 0.0206 |
| Block | 1, 17938 | 6.64 | 0.0099 |
| Trial Length | 2, 17938 | 51.31 | < 0.0001 |
| Hemisphere | 1, 52 | 2.48 | 0.1213 |
| Block*Trial Length | 2, 17938 | 11.66 | < 0.0001 |
| Block*Hemisphere | 1, 17938 | 3.31 | 0.0689 |
| Trial Length*Hemisphere | 2, 17938 | 1.98 | 0.1386 |
| Block*Trial Length*Hemisphere | 2, 17938 | 3.43 | 0.0325 |

**Supplemental Table 2. Estimated marginal means (EMM) and standard errors (SE) for inclusion trial lengths across blocks.** Asterisks (*) indicate EMMs that are significantly greater than zero and significant differences between different trial lengths within each block and between blocks. The effect column describes the direction of significant effects. ****p* < 0.0001, ***p* < 0.005, **p* < 0.05

| **Hemisphere** | **Trial Length** | **Fair Play Block**  **EMM (SE)** | **Ostracism Block**  **EMM( SE)** | **Trial Effect Within Block** | **Trial Effect Across Block** |
| --- | --- | --- | --- | --- | --- |
| **Left** | 2 s | 0.70 (0.21)** | -1.26 (0.50)* | Fair play 2 s > Fair play 4 s***  Fair play 2 s > Fair play 8 s**  Ostracism 2 s > Ostracism 8 s***  Ostracism 4 s > Ostracism 8 s*** | Fair play 2 s > Ostracism 2 s**  Fair play 8 s > Ostracism 8 s*** |
|  | 4 s | -0.86 (0.19)** | -1.76 (0.53)** |  |  |
|  | 8 s | -0.56 (0.27) | -4.53 (0.68)*** |  |  |
| **Right** | 2 s | 0.53 (0.21)* | -0.17 (0.50) | Fair play 2 s > Fair play 4 s***  Ostracism 2 s > Ostracism 4 s**  Ostracism 2 s > Ostracism 8 s** | Fair play 8 s > Ostracism 8 s* |
|  | 4 s | -0.70 (0.19)** | -1.86 (0.53)** |  |  |
|  | 8 s | -0.04 (0.27) | -2.55 (0.67)** |  |  |

**Supplemental Table 3. Model estimates for manipulation check using each Need-Threat Survey (NTS) outcome.** Estimated marginal means (EMM) and standard errors (SE) for each block are also reported.

| **Outcome** | **Term** | **DF** | ***F*-value** | ***p*-value** | **Fair-Play**  **EMM (SE)** | **Ostracism**  **EMM (SE)** |
| --- | --- | --- | --- | --- | --- | --- |
| **Estimated % of Throws Received** | Intercept | 1, 50 | 450.26 | <0.0001 | 24.90 (1.05) | 10.00 (1.01) |
|  | Block | 1, 46 | 133.76 | <0.0001 |  |  |
| **Feeling of Exclusion** | Intercept | 1, 52 | 731.11 | <0.0001 | 2.15 (0.14) | 4.42 (0.14) |
|  | Block | 1, 52 | 208.38 | <0.0001 |  |  |
| **Threat to Belonging** | Intercept | 1, 52 | 1452.31 | <0.0001 | 2.52 (0.11) | 4.18 (0.11) |
|  | Block | 1, 52 | 199.00 | <0.0001 |  |  |
| **Threat to Control** | Intercept | 1, 52 | 2152.50 | <0.0001 | 3.49 (0.10) | 4.47 (0.10) |
|  | Block | 1, 52 | 97.30 | <0.0001 |  |  |
| **Threat to Meaningful Existence** | Intercept | 1, 52 | 823.80 | <0.0001 | 2.38 (0.13) | 3.90 (0.13) |
|  | Block | 1, 52 | 141.04 | <0.0001 |  |  |
| **Threat to Self-Esteem** | Intercept | 1, 52 | 861.97 | <0.0001 | 2.52 (0.12) | 3.75 (0.12) |
|  | Block | 1, 52 | 114.65 | <0.0001 |  |  |
| **Positive Affect** | Intercept | 1, 52 | 652.43 | <0.0001 | 3.32 (0.12) | 2.05 (0.12) |
|  | Block | 1, 52 | 105.52 | <0.0001 |  |  |
| **Negative Affect** | Intercept | 1, 52 | 445.22 | <0.0001 | 1.54 (0.12) | 2.96 (0.12) |
|  | Block | 1, 52 | 146.10 | <0.0001 |  |  |

**Supplemental Table 4. Model estimates predicting effects of block, social threat, and social anxiety on z-scored Δ HbO.** LSAS = Liebowitz Social Anxiety Scale

|  | **Main Effect** | | | **Interaction with Block** | | **Interaction with Hemisphere** | | **Interaction with Block*Hemisphere** | |
| --- | --- | --- | --- | --- | --- | --- | --- | --- | --- |
|  | **DF** | ***F*-value** | ***p*-value** | ***F*-value** | ***p*-value** | ***F*-value** | ***p*-value** | ***F*-value** | ***p*-value** |
| **Intercept** | 1, 15621 | 4.37 | **0.0366** | - | - | - | - | - | - |
| **Block** | 1, 15621 | 8.90 | **0.0029** | - | - | - | - | - | - |
| **Hemisphere** | 1, 51 | 0.01 | 0.9382 | 0.74 | 0.3888 | - | - | - | - |
| **Threat to Belonging** | 1, 15621 | 0.79 | 0.3730 | 0.21 | 0.6430 | 0.00 | 0.9969 | 0.02 | 0.8955 |
| **Threat to Control** | 1, 15621 | 0.90 | 0.3436 | 1.04 | 0.3073 | 0.28 | 0.5958 | 0.85 | 0.3565 |
| **Threat to Meaningful Existence** | 1, 15621 | 0.88 | 0.3471 | 0.00 | 0.9716 | 0.81 | 0.3686 | 5.22 | **0.0223** |
| **Threat to Self-Esteem** | 1, 15621 | 0.14 | 0.7034 | 0.16 | 0.6875 | 0.00 | 0.9647 | 0.80 | 0.3698 |
| **LSAS** | 1, 51 | 2.50 | 0.1200 | 0.71 | 0.3986 | 0.05 | 0.8218 | 2.01 | 0.1559 |
| **Threat to Belonging*LSAS** | 1, 15621 | 0.02 | 0.8777 | 0.30 | 0.5820 | 0.33 | 0.5631 | 1.66 | 0.1972 |
| **Threat to Control*LSAS** | 1, 15621 | 8.14 | **0.0043** | 0.00 | 0.9678 | 0.38 | 0.5392 | 0.02 | 0.8833 |
| **Threat to Meaningful Existence*LSAS** | 1, 15621 | 0.42 | 0.5162 | 0.50 | 0.4795 | 1.90 | 0.1677 | 0.34 | 0.5611 |
| **Threat to Self-Esteem*LSAS** | 1, 15621 | 3.38 | 0.0661 | 2.83 | 0.0927 | 0.69 | 0.4057 | 1.15 | 0.2826 |

**Supplemental Table 5. Model estimates predicting effects of block, social threat, and need for belonging on z-scored Δ HbO.** NTBS = Need to Belong Scale

|  | **Main Effect** | | | **Interaction with Block** | | **Interaction with Hemisphere** | | **Interaction with Block*Hemisphere** | |
| --- | --- | --- | --- | --- | --- | --- | --- | --- | --- |
|  | **DF** | ***F*-value** | ***p*-value** | ***F*-value** | ***p*-value** | ***F*-value** | ***p*-value** | ***F*-value** | ***p*-value** |
| **Intercept** | 1, 15621 | 4.76 | **0.0291** | - | - | - | - | - | - |
| **Block** | 1, 15621 | 7.69 | **0.0055** | - | - | - | - | - | - |
| **Hemisphere** | 1, 51 | 0.02 | 0.8979 | 0.82 | 0.3663 | - | - | - | - |
| **Threat to Belonging** | 1, 15621 | 0.63 | 0.4259 | 0.29 | 0.5913 | 0.00 | 0.9963 | 0.02 | 0.8811 |
| **Threat to Control** | 1, 15621 | 0.51 | 0.4751 | 0.41 | 0.5226 | 0.53 | 0.4656 | 1.30 | 0.2539 |
| **Threat to Meaningful Existence** | 1, 15621 | 0.89 | 0.3453 | 0.08 | 0.7752 | 1.10 | 0.2933 | 3.94 | **0.0472** |
| **Threat to Self-Esteem** | 1, 15621 | 0.54 | 0.4611 | 0.10 | 0.7491 | 0.00 | 0.9663 | 1.61 | 0.2044 |
| **NTBS** | 1, 51 | 0.01 | 0.9168 | 0.28 | 0.5977 | 0.04 | 0.8353 | 1.60 | 0.21 |
| **Threat to Belonging*NTBS** | 1, 15621 | 2.12 | 0.1458 | 0.65 | 0.4204 | 0.08 | 0.7810 | 0.45 | 0.5035 |
| **Threat to Control*NTBS** | 1, 15621 | 3.02 | 0.0823 | 2.76 | 0.0968 | 1.85 | 0.1733 | 4.20 | **0.0405** |
| **Threat to Meaningful Existence*NTBS** | 1, 15621 | 0.00 | 0.9986 | 0.02 | 0.8770 | 0.63 | 0.4283 | 0.03 | 0.8669 |
| **Threat to Self-Esteem*NTBS** | 1, 15621 | 0.12 | 0.7260 | 0.20 | 0.6543 | 0.18 | 0.6722 | 0.88 | 0.3483 |

**Supplemental Table 6. Model estimates predicting effects of block, social threat, and personality growth mindset on z-scored Δ HbO.** ITP = Implicit Theories of Personality

|  | **Main Effect** | | | **Interaction with Block** | | **Interaction with Hemisphere** | | **Interaction with Block*Hemisphere** | |
| --- | --- | --- | --- | --- | --- | --- | --- | --- | --- |
|  | **DF** | ***F*-value** | ***p*-value** | ***F*-value** | ***p*-value** | ***F*-value** | ***p*-value** | ***F*-value** | ***p*-value** |
| **Intercept** | 1, 15621 | 4.08 | **0.0434** | - | - | - | - | - | - |
| **Block** | 1, 15621 | 7.09 | **0.0078** | - | - | - | - | - | - |
| **Hemisphere** | 1, 51 | 0.01 | 0.9047 | 0.80 | 0.3695 | - | - | - | - |
| **Threat to Belonging** | 1, 15621 | 0.58 | 0.4470 | 0.21 | 0.6477 | 0.00 | 0.9964 | 0.00 | 0.9398 |
| **Threat to Control** | 1, 15621 | 0.33 | 0.5644 | 0.28 | 0.5942 | 0.56 | 0.4553 | 0.59 | 0.4427 |
| **Threat to Meaningful Existence** | 1, 15621 | 1.19 | 0.2751 | 0.27 | 0.6026 | 0.84 | 0.3577 | 3.67 | 0.0555 |
| **Threat to Self-Esteem** | 1, 15621 | 0.45 | 0.5021 | 0.00 | 0.9501 | 0.00 | 0.9440 | 1.29 | 0.2553 |
| **ITP** | 1, 51 | 0.35 | 0.5575 | 0.21 | 0.6461 | 1.12 | 0.2945 | 0.05 | 0.8222 |
| **Threat to Belonging*ITP** | 1, 15621 | 2.11 | 0.1459 | 0.49 | 0.4815 | 0.02 | 0.8750 | 3.57 | 0.0587 |
| **Threat to Control*ITP** | 1, 15621 | 0.16 | 0.6927 | 0.47 | 0.4915 | 0.00 | 0.9675 | 0.23 | 0.6279 |
| **Threat to Meaningful Existence*ITP** | 1, 15621 | 4.07 | **0.0437** | 0.32 | 0.5735 | 0.32 | 0.5681 | 2.50 | 0.1135 |
| **Threat to Self-Esteem*ITP** | 1, 15621 | 0.00 | 0.9520 | 0.00 | 0.9872 | 0.27 | 0.6012 | 0.33 | 0.5653 |

**Supplemental Figure 1. Violin plots of z-scored ΔHbO concentration across blocks and hemispheres.** Across both panels, individual points and lines connecting points indicate individual participants over the two blocks. Turquoise represents the fair play block and purple represents the ostracism block. **Left panel** represents differences between left and right vlPFC activity for the fair play block. **Right panel** represents differences between left and right vlPFC activity for the ostracism block.

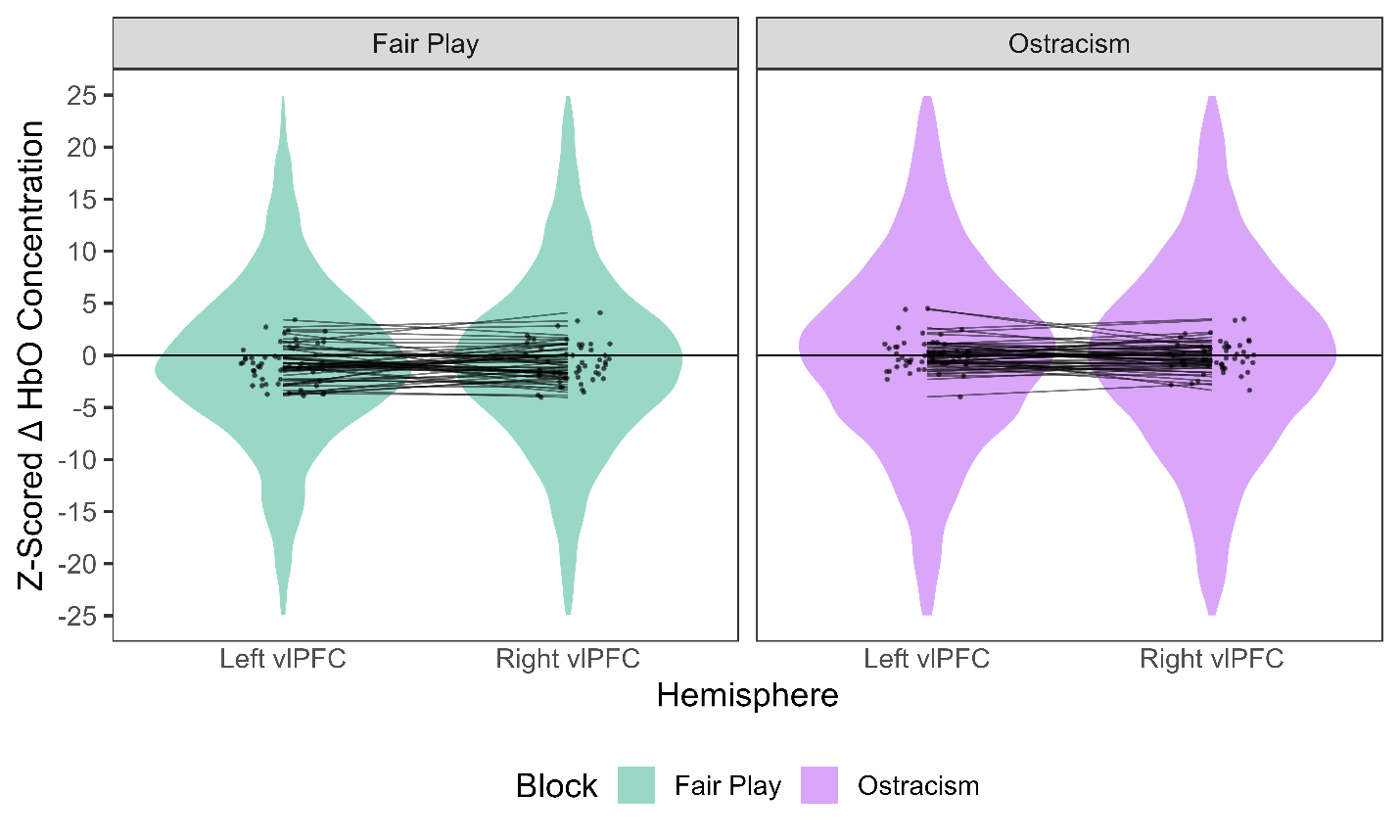
